# INDEHISCENT regulates explosive seed dispersal

**DOI:** 10.1101/2021.06.11.448014

**Authors:** Anahit Galstyan, Penny Sarchet, Rafael Campos-Martin, Milad Adibi, Lachezar A. Nikolov, Miguel Pérez Antón, Léa Rambaud-Lavigne, Xiangchao Gan, Angela Hay

## Abstract

How traits evolve to produce novelty or stasis is an open question in biology. We investigate this question in *Cardamine hirsuta*, a relative of *Arabidopsis thaliana* that employs explosive fracture to disperse its seeds. This trait evolved through key morphomechanical innovations that distinguish the otherwise very similar, dehiscent fruit of these two species. Using CRISPR/Cas9, we show that dehiscence zone formation is absolutely required for explosive fracture in *C. hirsuta*, and is controlled by the bHLH transcription factor INDEHISCENT (IND). Using mutant screens, we identified the MADS-box transcription factor FRUITFULL (FUL) as a negative regulator of *IND* in *C. hirsuta*. Although FUL function is conserved in *C. hirsuta*, the consequences of *IND* mis-expression differ in *ful* mutants of *C. hirsuta* versus *A. thaliana*. In *ful* mutants of both species, valve tissue is replaced by dehiscence zone tissue, which comprises two distinct cell types: lignified layer and separation layer cells. While *A. thaliana ful* mutants develop stunted fruit with ectopic lignified layer cells, *C. hirsuta ful* mutants have elongated fruit with ectopic separation layer cells. We show that IND dose determines the proportion of these two cell types in ectopic dehiscence zones. We also show that the extent of ectopic lignification caused by *IND* mis-expression determines fruit length. Our findings indicate developmental system drift in the conserved gene network patterning dehiscent fruit in two related Brassicaceae species.

## Introduction

Adaptations for dispersal are ubiquitous in nature. By moving individuals from one area to another, dispersal has important ecological and evolutionary consequences – including the ability to change or expand a species’ range, which is especially relevant in today’s context of climate change and landscape fragmentation (Kokko and Lopez-Sepulcre, 2006). *Cardamine hirsuta* uses an explosive mechanism to disperse its seeds in a wide radius around the parent plant (Hofhuis *et al*., 2016). While various flowering plants employ this strategy of explosive seed dispersal, it is restricted to *Cardamine* within the Brassicaceae family, and therefore absent in the model species *Arabidopsis thaliana.* We took advantage of this difference in seed dispersal strategy between two closely related species, to investigate the degree of conservation versus divergence in the gene regulatory networks that pattern explosive versus non-explosive fruit tissues.

In *C. hirsuta*, explosive fracture converts elastic potential energy in the fruit valves to kinetic coiling energy to eject the seeds (Hofhuis *et al*., 2016). The fracture runs along the seam that connects the coiling valve to the fruit. This connecting seam is called the dehiscence zone. The dehiscent fruit of both *C. hirsuta* and *A. thaliana* comprise two valves that are joined margin to margin along the replum and enclose the seeds. At maturity, the valves separate from the fruit along the dehiscence zone. In *A. thaliana*, this process occurs as the fruit dries, leaving the seeds exposed for subsequent dispersal by external agents (Dinneny and Yanofsky, 2005). So, the developmentally controlled process of dehiscence is temporally distinct from seed dispersal. In *C. hirsuta*, by contrast, dehiscence coincides with seed dispersal when the fruit explodes before drying. Therefore, the genes that pattern dehiscence zone formation are likely to influence explosive seed dispersal in *C. hirsuta*.

The patterning mechanism that draws a line of cells with dehiscence zone fate along the valve margins, has been elucidated from genetic studies in *A. thaliana* (Dinneny and Yanofsky, 2005). The spatial precision of this patterning is controlled by a group of genes that promote dehiscence zone cell fate at the valve margins, and another group of genes that restrict this fate in surrounding tissues. Four genes encode transcription factors that promote dehiscence zone fate: the MADS-box proteins SHATTERPROOF 1 (SHP1) and SHP2, and the basic helix-loop-helix (bHLH) proteins INDEHISCENT (IND) and ALCATRAZ (ALC) (Liljegren *et al*., 2000; Liljegren *et al*., 2004; Rajani and Sundaresan, 2001). Expression of these four genes is mostly limited to the valve margins through repression by the MADS-box transcription factor FRUITFULL (FUL) in the valves and by the homeodomain transcription factor REPLUMLESS in the replum (Ferrandiz *et al*., 2000b; Liljegren *et al*., 2004; Roeder *et al*., 2003). This patterning mechanism ensures that three tissues differentiate post-fertilization in the fruit – valve, replum and dehiscence zone -with sharp borders between them.

Successful dehiscence depends on the differentiation of two specialized cell types in the dehiscence zone. A layer of lignified cells forms adjacent to the valve and a layer of non-lignified cells, capable of autolysis, forms adjacent to the replum (Spence *et al*., 1996). Lignified cell fate and complete dehiscence require the transcription factor NAC SECONDARY WALL THICKENING PROMOTING FACTOR 1 (Mitsuda and Ohme-Takagi, 2008). While the non-lignified cells require the enzymatic activity of ARABIDOPSIS DEHISCENCE ZONE POLYGALACTURONASE 1 (ADPG1) and ADPG2 to degrade cell wall pectins, causing cells to separate and break (Ogawa et al., 2009). In this way, the valve separates from the replum along a precise cell layer at the valve margin.

The key regulator of dehiscence zone cell fate in *A. thaliana* is IND (Liljegren *et al*., 2004). Loss of *IND* function causes loss of both the lignified layer and separation layer, while *ALC* is required only for separation layer fate (Liljegren *et al*., 2004; Rajani and Sundaresan, 2001). The expression of both *IND* and *ALC* is activated by SHP1/2 and these two target genes account for the majority of SHP1/2 function in valve margin development (Liljegren *et al*., 2000; Liljegren *et al*., 2004). However, each gene has distinct as well as overlapping functions, suggesting they form a nonlinear regulatory network (Liljegren *et al*., 2004). For example, IND forms heterodimers with the related bHLH proteins ALC and SPATULA (SPT), and ALC and SPT can also heterodimerise with each other and interact with DELLA repressor proteins (Arnaud *et al*., 2010; Gallego-Bartolome *et al*., 2010; Girin *et al*., 2011; Groszmann *et al*., 2011; Liljegren *et al*., 2004). The gibberellin (GA) biosynthesis gene *GA3ox1* and *SPT* are direct targets of IND (Arnaud *et al*., 2010; Girin *et al*., 2011). Therefore, local GA production in the valve margin leads to DELLA protein degradation and release of ALC, and likely SPT (Arnaud *et al*., 2010; Gallego-Bartolome *et al*., 2010). In this way, IND contributes to the activation of *ALC* and *SPT* in the valve margin, where they regulate genes that cause cell separation along the dehiscence zone. An additional function of ALC and SPT is to feedback and repress *IND* gene expression (Lenser and Theissen, 2013). Although cellular resolution for these regulatory interactions is currently missing, this interplay between IND, DELLAs and ALC/SPT may provide a way to delimit the differentiation of separation and lignified layers in the dehiscence zone (Ballester and Ferrandiz, 2017).

Transcriptional targets of IND also include the AGC3 kinase genes *PINOID* (*PID*) and *WAG2* (Girin *et al*., 2011; Sorefan *et al*., 2009). Manipulating auxin levels in the *IND* expression domain - for example, increasing auxin synthesis by expressing the bacterial *iaaM* gene, or redistributing auxin by expressing *PID* or *WAG2* – results in indehiscent fruit (Sorefan *et al*., 2009; van Gelderen *et al*., 2016). PID and WAG2 regulate abundance of the PIN-FORMED 3 auxin efflux carrier at the plasma membrane, and, in this way, IND controls auxin distribution at the valve margin (Sorefan *et a*l., 2009; van Gelderen *et al*., 2016). However, auxin responses are dynamic throughout valve margin development, with an initial maximum during dehiscence zone formation transitioning to a minimum during the differentiation of dehiscence zone cells (Li *et al*., 2019; Sorefan *et al*., 2009; van Gelderen *et al*., 2016). Therefore, it is still not clear exactly when and how IND regulates auxin to ensure successful dehiscence (Ballester and Ferrandiz, 2017).

IND appears to be a conserved regulator of fruit dehiscence within the Brassicaceae. IND is an atypical bHLH protein, found only in the Brassicaceae, and reverse genetics in *Brassica rapa*, *Brassica oleracea*, *Capsella rubella* and *Lepidium campestre* have shown that loss of *IND* function results in indehiscent fruit (Bailey *et al*., 2003; Dong *et al*., 2019; Girin *et al*., 2010; Heim *et al*., 2003; Kay *et al*., 2013; Lenser and Theissen, 2013; Toledo-Ortiz *et al*., 2003). *IND* expression in the valve margin of *A. thaliana* is governed by a 406-bp promoter sequence, and this conserved non-coding region has been identified in eight other Brassicaceae *IND* genes (Dong *et al*., 2019; Girin *et al*., 2010). In *Capsella rubella*, regulatory sequences outside of this region expand the domain of *IND* expression into the valves and partially contribute to the heart-shaped form of these fruit (Dong *et al*., 2019). Despite this derived expression domain, the function of FUL to repress *IND* expression is conserved in *C. rubella* (Dong *et al*., 2019). Given these examples, it is important to investigate the degree of conservation versus divergence in the gene regulatory networks that pattern diverse fruit within the Brassicaceae.

By taking a genetic approach in *Cardamine hirsuta*, we show that dehiscence zone formation is absolutely required for explosive fracture of the fruit. In this way, developmental regulators of dehiscence zone formation control explosive seed dispersal in *C. hirsuta*. We show that *IND* and *FUL* have conserved roles in patterning the dehiscent fruit of *C. hirsuta*. In *ful* mutants, dehiscence zones replace valves as a result of ectopic *IND* expression. However, this ectopic tissue comprises mostly separation layer cells in *C. hirsuta ful*, compared with lignified layer cells in *A. thaliana ful*. We show that IND dose determines the proportion of these two dehiscence zone cell types, suggesting that quantitative changes affecting *IND* expression in this conserved gene network may explain developmental system drift between two closely related species.

## Results

### Dehiscence zone formation in *C. hirsuta*

Dehiscence zones form in *C. hirsuta* fruit at the valve margins, adjacent to the replum, and comprise lignified and separation cell layers, similar to the non-explosive fruit of *A. thaliana* (Fig. 1A-D). Dehiscence zones form at the valve margins during early stages of fruit development (14–16, Fig. 1E-K) and differentiate lignified and separation layer cell types during later stages (17a–17b, Fig. L-O). A narrow band of cells at the valve margin begin to divide at stage 14 to produce a region of smaller cells at the valve margin (Fig. 1F-K). The fruit surface starts to indent at this region of smaller cells between the replum and valve at stage 15 (Fig. 1G). Cell divisions are no longer observed after stage 16 (Fig. 1K-O), suggesting that further growth of the dehiscence zone occurs via cell expansion.

**Figure 1.**
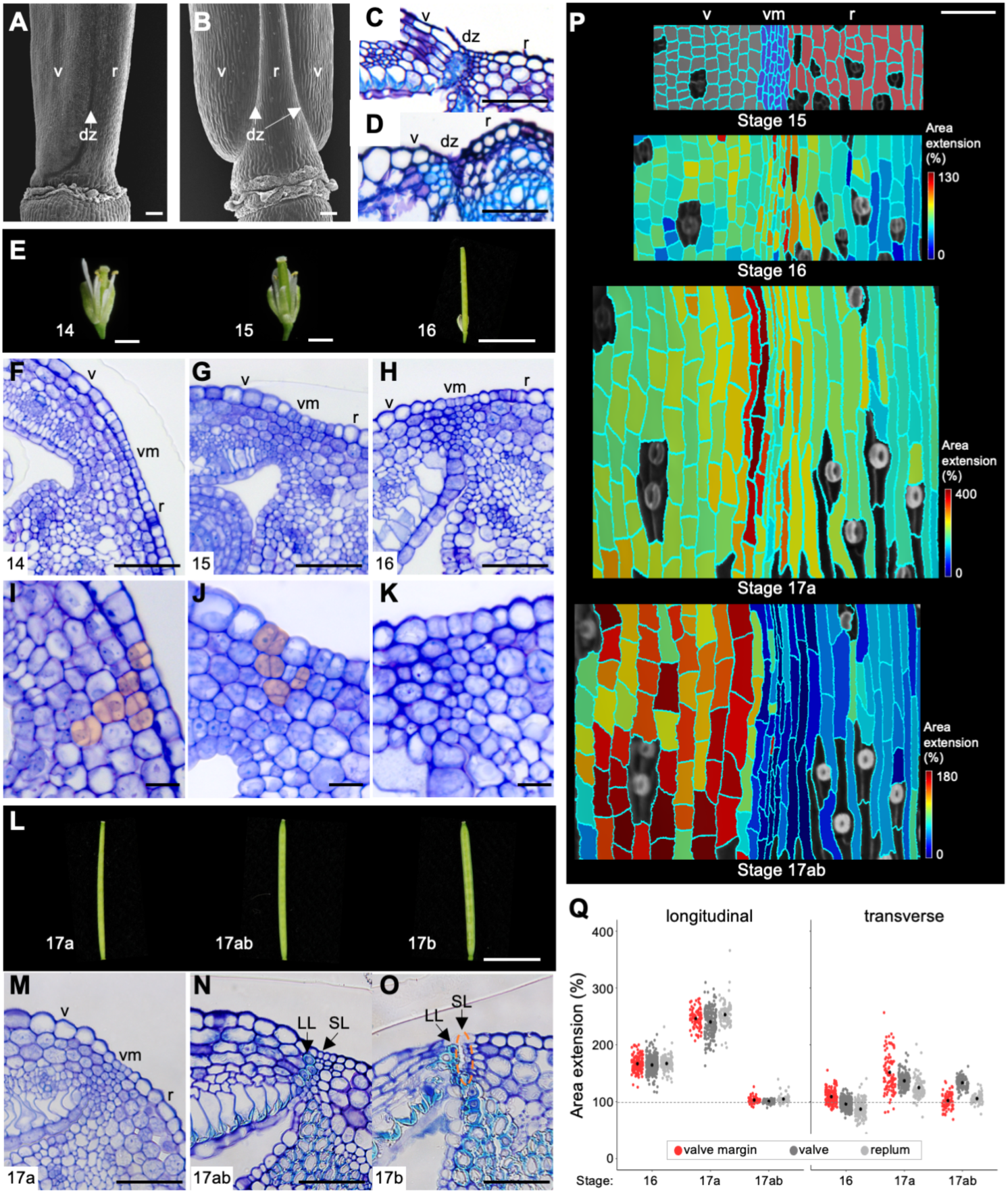
Dehiscence zone development in *C. hirsuta* fruit. (A-D) *C. hirsuta* (A, C) and *A. thaliana* (B, D) stage 17b fruit base in SEM (A-B) and TBOstained transverse sections of the dehiscence zone (C-D); lignified cell walls stain cyan. (E-K) *C. hirsuta* stage 14 to 16 fruit (E) and TBO-stained transverse sections of the valve margin in stage 14 (F,I), 15 (G, J), 16 (H, K) fruit; close-up views of the valve margin with recently divided cells indicated in orange (I-J). (L-O) *C. hirsuta* stage 17a to 17b fruit (L) and TBO-stained transverse sections of the valve margin in stage 17a (M), 17ab (N), 17b (O) fruit; lignified cell walls stain cyan; dashed orange circle indicates separated cells. (P-Q) Cellular growth in a time-lapse series of *C. hirsuta* fruit during stages 15 to 17ab. Heat maps indicate cell area extension (%) between stages, each heat map scales the data between maximum and minimum values; fruit tissue types labelled at stage 15 (valve: grey, valve margin: blue, replum: red) (P). Cell area extension (%) between stages in the longitudinal and transverse directions of the fruit, in valve margin (red), valve (dark grey) and replum (light grey); black dots indicate means; dashed line indicates no change in cell area; n = 1308 cells (Q). Abbreviations: v, valve; r, replum; dz, dehiscence zone; vm, valve margin; LL, lignified layer; SL, separation layer. Scale bars: 100 μm (A, B), 50 μm (C, D, F-H, M-P), 10 μm (I-K).

To quantify cellular growth during dehiscence zone formation, we used live confocal imaging. We imaged epidermal cell outlines at four stages over seven days of fruit development and quantified cell area extension using MorphoGraphX in two replicate fruit series (Fig. 1P, Fig. S1) (Barbier de Reuille *et al*., 2015). We observed no cell division in these samples, indicating that fruit epidermal cells grew by expansion during this time. Growth was distributed across all three tissues of the fruit up until stage 17a, with maximal growth occurring between stages 16 and 17a (Fig. 1P). Cell area extension was higher in the longitudinal direction of the fruit in all tissues up to stage 17a (Fig. 1Q), resulting in elongation of the fruit to its maximal length. Growth ceased after stage 17a in all tissues except for the valve, which continued to grow in the transverse direction of the fruit (Fig. 1P, Q), contributing to the expansion of fruit width. In summary, cells in the valve margin are significantly smaller than valve or replum cells (Fig. S2). They have a thin shape and grow predominantly in the longitudinal direction of the fruit, resulting in small, thin cells at the valve margin (Fig. 1Q, Fig. S2).

Cell differentiation of the dehiscence zone occurs during stage 17ab to form a lignified cell layer, adjacent to the valve, and a separation cell layer, adjacent to the replum (arrows, Fig. 1N). At this stage, lignification also occurs in endocarp *b* cells in the valve and lignified cell types in the replum (Fig. 1N). The separation layer can be identified as a file of small cells flanked by lignified layer cells and larger replum cells (arrow, Fig. 1N). Cells along the separation layer physically detach from the lignified cell layer during stage 17b (dashed circle, Fig. 1O). This complete maturation of the dehiscence zone coincides with when *C. hirsuta* valves acquire the capacity to explosively coil.

### Dehiscence zone is required for explosive pod shatter

Given the spatial and temporal association between differentiation of the dehiscence zone and explosive fracture along this tissue, we tested whether dehiscence is required for explosive pod shatter. To generate *C. hirsuta* fruit that lack dehiscence zones, we performed CRISPR/Cas9 gene editing of *C. hirsuta IND*. We generated a *C. hirsuta ind-1* allele with a single nucleotide insertion at position 361 bp of the coding sequence (Fig. S3). This insertion introduces a frame shift at amino acid position 121 and a premature stop codon, producing a truncated protein of 182 amino acids with 62 amino acids of nonsense protein sequence at the C-terminus (Fig. S3). This truncated protein lacks the basic helix-loop-helix domain that is critical for DNA binding and protein dimerization (Carretero-Paulet *et al*., 2010), suggesting that *C. hirsuta ind-1* is a loss-of-function allele (Fig. S3). Plants homozygous for the recessive *C. hirsuta ind-1* allele produced non-explosive, indehiscent fruit (Fig. 2A-B). These valves can coil when manually detached from the replum, but fail to eject the seeds (Fig. 2B). To demonstrate that loss of explosive seed dispersal in *C. hirsuta ind-1* is caused by loss of *IND* function, we fully complemented the mutant phenotype with a *pChIND::ChIND:VENUS* transgene (Fig. 2I-K). Therefore, *C. hirsuta IND* is required for explosive pod shatter.

**Figure 2.**
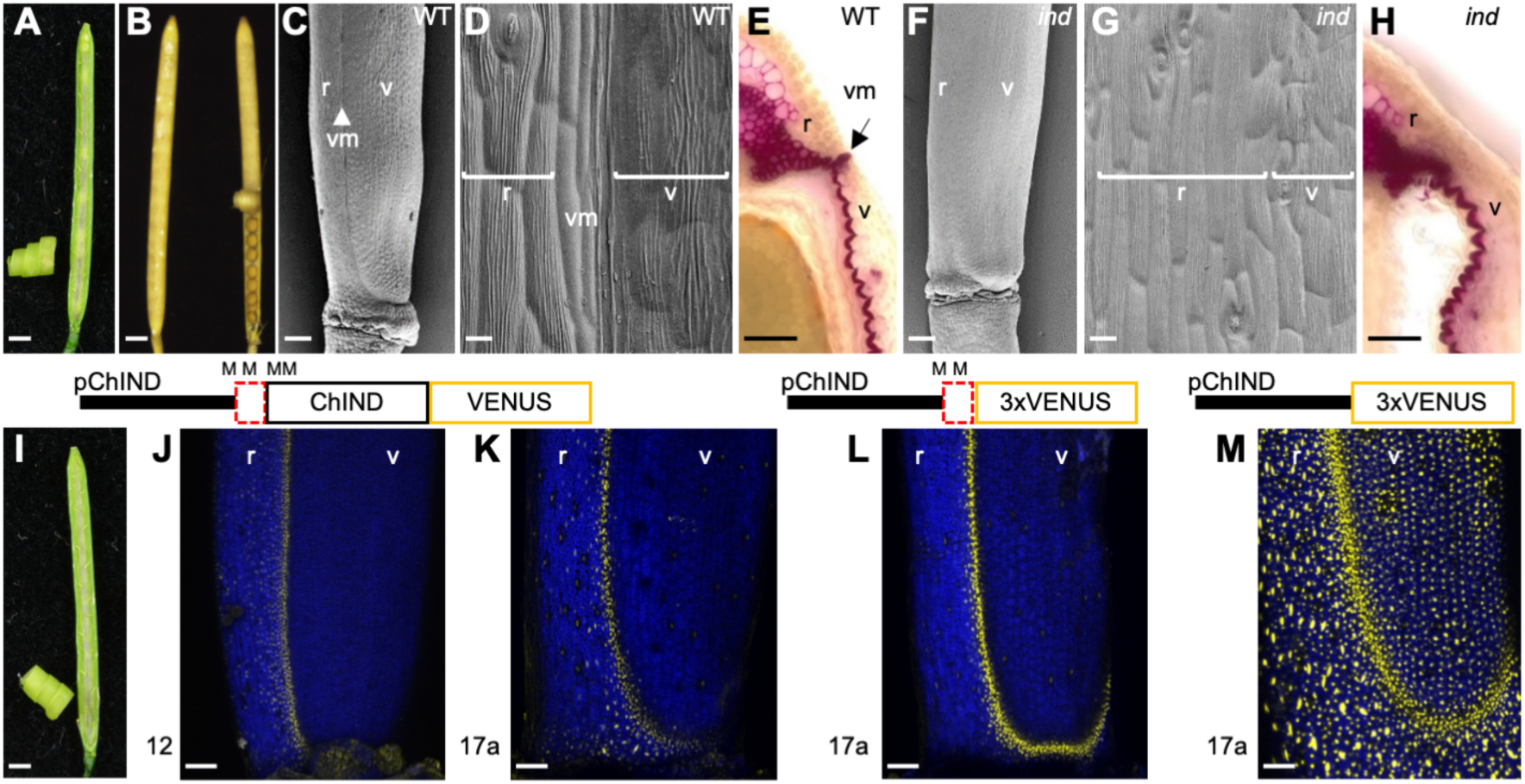
*C. hirsuta ind* mutant has non-explosive, indehiscent fruit. (A-B) *C. hirsuta* stage 17b wild-type dehisced fruit and coiled valve (A), stage 18 *ind-1* indehiscent fruit with coiled valve manually detached on right (B). (C-H) SEM (C-D, F-G) and phloroglucinol stained transverse sections (E, H) of stage 17ab *C. hirsuta* wild type and *ind-1* fruit; lignified cell walls stain pink. (I-K) Fruit of *ind-1* complemented with *pChIND::ChIND:VENUS* transgene (I), and CLSM of pChIND::ChIND:VENUS expression in stage 12 (J) and stage 17a (K) *ind-1* fruit; n = 11 independent single insertion T2 lines. (L-M) CLSM of pChIND-UTR::3xVENUS (L) and pChIND::3xVENUS (M) transcriptional reporters in wild type stage 17a fruit; n = 6 independent T1 lines. Cartoons of each construct are shown above panels; M indicates predicted Met codons. Venus fluorescence (yellow), chlorophyll autofluorescence (blue). Scale bars: 1mm (A, B, I), 200μm (C, F), 10μm (D, G) 50μm (E, H, J-M). Abbreviations: v, valve; r, replum; vm, valve margin.

The indehiscent fruit of *C. hirsuta ind-1* completely lack the narrow band of valve margin tissue, characterized in wild-type fruit by thin cells and no stomata (Fig. 2C, D, F, G). Using phloroglucinol to specifically stain lignin, we found no lignified cells at the valve margin of *C. hirsuta ind-1* fruit (Fig. 2E, H). However, epidermal features and lignification patterns of valve and replum tissues appeared normal in mutant fruit (Fig. 2F-H). Therefore, *IND* functions as a master regulator of dehiscence zone formation in *C. hirsute* as it does in *A. thaliana*.

To analyze the accumulation of *C. hirsuta* IND protein during fruit development, we imaged the *pChIND::ChIND:VENUS* transgenic lines. A sharp boundary is established early in fruit development between IND expression in the valve margin and its absence from the rest of the valve (stage 12, Fig. 2J). This sharp boundary is maintained throughout subsequent stages of fruit development with strong expression of IND in the valve margins (stages 17a, Fig. 2K). In contrast to the valve, we observed IND expression extending from the valve margin part way into the replum (Fig. 2J-K). To map the DNA sequences required for tissue-specific expression of *C. hirsuta* IND, we generated transcriptional fusions using either 1885 bp of non-transcribed promoter sequence (*pChIND::3xVENUS*), which includes the conserved non-coding sequence that is sufficient for valve margin expression of *A. thaliana IND* (Girin *et al*., 2010) (Fig. S3), or an additional 84 bp of transcribed 5’ UTR (*pChIND-UTR::3xVENUS*) (Fig. 2L-M, Fig. S3). We found that valve margin-specific expression of *C. hirsuta IND* strictly requires the 84 bp 5’UTR (Fig. 2L), as the 1885 bp promoter fusion was expressed ubiquitously throughout the fruit (Fig. 2M). In summary, *C. hirsuta* IND accumulates in the valve margin prior to development of the dehiscence zone and is maintained throughout its differentiation.

### Genetic screen for indehiscent mutants in *C. hirsuta*

To identify additional regulators of dehiscence zone formation in *C. hirsuta*, we screened ethyl methanesulfonate-mutagenized plants for mutants with indehiscent fruit. We named an indehiscent mutant from this screen *valveless* because its fruit lacked valves (Fig. 3A-B). Using map-based cloning, we identified a C to T nucleotide substitution in the *C. hirsuta* MADS-box gene *FRUITFULL* (*FUL*, gene identifier CARHR274590 (Gan *et al*., 2016)) that co-segregated with the *valveless* mutant phenotype (Fig. S4). This mutation introduces a premature stop codon that results in a truncated 143 amino acid protein that contains the MADS-box but not the K-box dimerization domain (Fig. S4). *C. hirsuta FUL* transcripts accumulate to their highest levels in the valves during fruit development and are reduced to 5% of wild-type levels in mutant fruit (Fig. 3D-E). In addition, silencing *C. hirsuta FUL* expression using VIGS, produced indehiscent fruit that lacked valve tissue and phenocopied the *valveless* mutant (Fig. S4). Moreover, the *valveless* mutant phenotype was fully complemented by introducing a *pChFUL::ChFUL:GFP* transgene (Fig. 3C). Taken together, these results show that *valveless* is a recessive, loss-of-function allele of *C. hirsuta FUL*, so we renamed the allele *C. hirsuta ful-1*.

**Figure 3.**
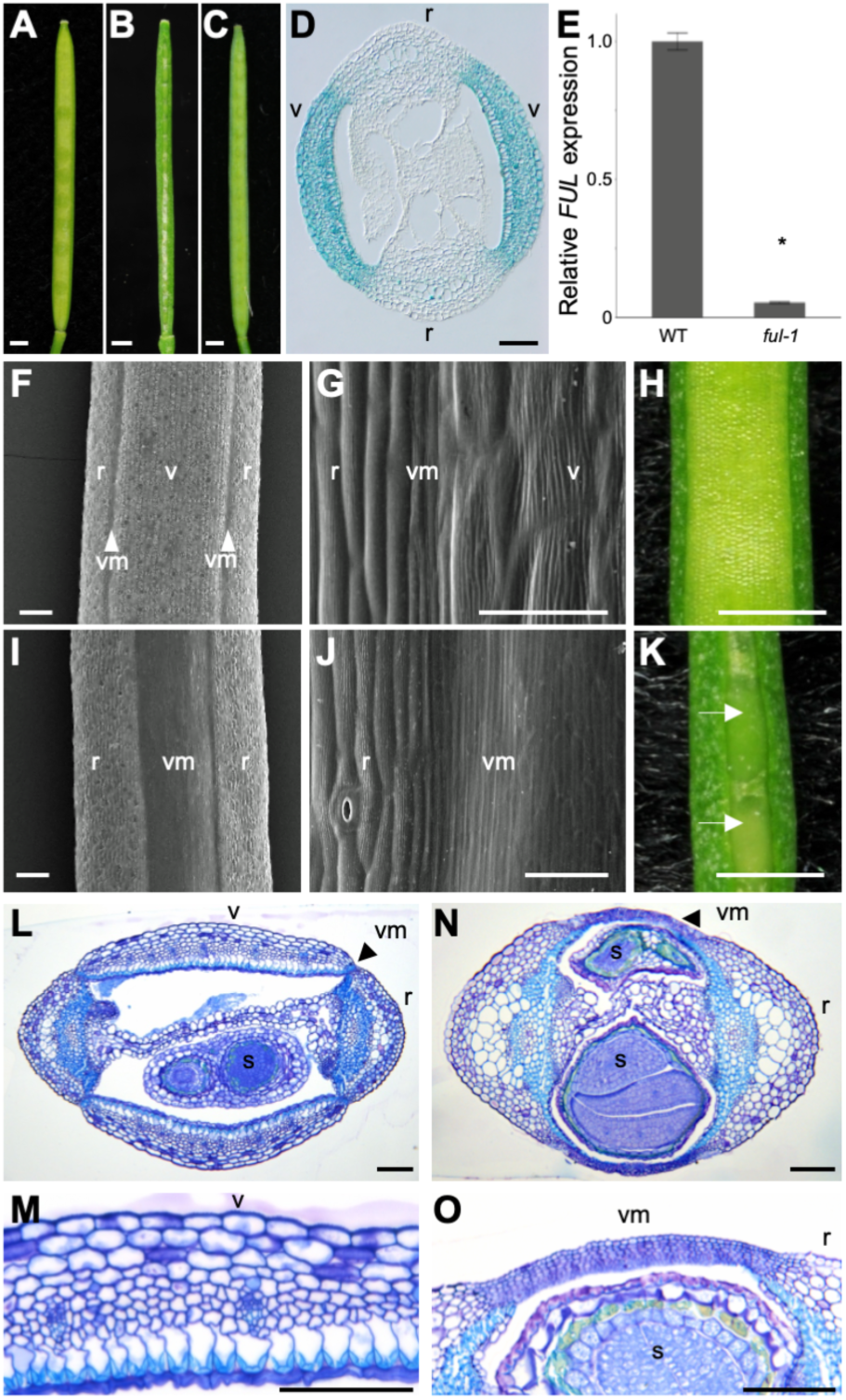
*C. hirsuta ful* mutant has indehiscent fruits that lack valves. (A-C) Fruit of *C. hirsuta* wild type (A), *ful-1* (B) and *ful-1* complemented with *pChFUL::ChFUL:VENUS* transgene (C), n=11 independent single insertion lines. (D) Transverse section of *pChFUL::GUS* fruit, blue colour indicates GUS expression in valves. (E) *C. hirsuta FUL* expression in wild-type and *ful-1* stage 17 fruit, determined by quantitative RT-PCR, normalized to expression of the housekeeping gene *CLATHRIN*, and shown relative to wild type, bars represent mean ±SEM of three biological replicates. Asterisk indicates statistically significant difference between wild type and *ful-1* (P <0.05; Studentˈs t-test). (F-K) Stage 17 fruit of wild type (F-H) and *ful-1* (I-K). SEMs compare epidermal cell types between wild-type (F-and *ful-1* (I-J) fruit. Arrows indicate visible seeds through the transparent tissue in *ful-1* fruit (K). (L-O) Toluidine Blue O-stained transverse sections of wild-type (L-M) and *ful-1* (N-O) fruit, close-up views of wild-type valve (M) and ectopic valve margin in *ful-1* (O); lignified cell walls stain cyan. Abbreviations: v, valve; r, replum; vm, valve margin; s, seed. Scale bar indicates 1 mm (A-C, H, K), 50 μm (D, G, J), 100 μm (F, I, L-O).

The fruit of *C. hirsuta ful-1* are comparable to wild type in length, but narrower in width (Fig. 3A-B). These mutant fruits lack valves, which are replaced by expanded regions of valve margin-like tissue (Fig. 3F-K). In wild-type fruit, the valve margin comprises thin cells with a smooth epidermal surface and no stomata (Fig. 3G). Valve cells have a striated epidermal surface, are wider than replum cells, and stomata are present in both valve and replum (Fig. 3F-G). Valve cells are absent from *C. hirsuta ful-1* fruit, while replum and valve margin-like cells are present (Fig. 3I-J). The ectopic valve margin-like tissue in the mutant is thin and semi-transparent such that seeds are clearly visible through it (Fig. 3K). Replum size is also expanded in *C. hirsuta ful-1* fruit, but despite this expansion in replum and valve margin tissues, the fruit remains narrow with very reduced locules in the absence of valves (Fig. 3L, N). As a consequence, the space available for seeds to develop is very constricted in *C. hirsuta ful-1* fruit, resulting in narrow, deformed seeds that are nonetheless fully viable (Fig. 3N, Fig. S5). We could not detect *FUL* transcripts in wild-type *C. hirsuta* seeds, and embryos developed normally in *C. hirsuta ful-1* mutants up until maturity when space restrictions caused some embryos to bend abnormally (Figs. S4, S5). Therefore, the primary defect in *C. hirsuta ful-1* fruit is a complete loss of valves and their replacement by valve margin-like tissue.

Comparison of wild-type valve with the valve margin-like tissue found in *C. hirsuta ful-1* fruit, shows an absence of differentiated valve cell types in the mutant (Fig. 3M, O). Wild-type valves comprise the exocarp cell layer, approximately 7 mesocarp cell layers, a characteristic endocarp *b* cell layer with asymmetric lignin deposition, and an endocarp *a* cell layer adjacent to the seeds (Fig. 3M). In comparison, the ectopic valve margin that forms in place of valve in *C. hirsuta ful-1*, comprises 9-10 layers of small cells that resemble the replum-adjacent separation layer of the wild-type dehiscence zone (Fig. 3O). Additionally, cells with uniform lignification, similar to lignified cells of the wild-type dehiscence zone, are sometimes found in the ectopic valve margin (Fig. 5C). These differences in histology between *C. hirsuta ful-1* and wild-type fruit are first observed at stage 12 and are clearly apparent by stage 14 when valve tissue starts to differentiate in wild type (Fig. S6). Taken together, our findings suggest that valve cell fate is not specified in *C. hirsuta ful-1* and instead, this fruit tissue adopts dehiscence zone cell fates, mostly of the replum-adjacent separation layer.

**Figure 4.**
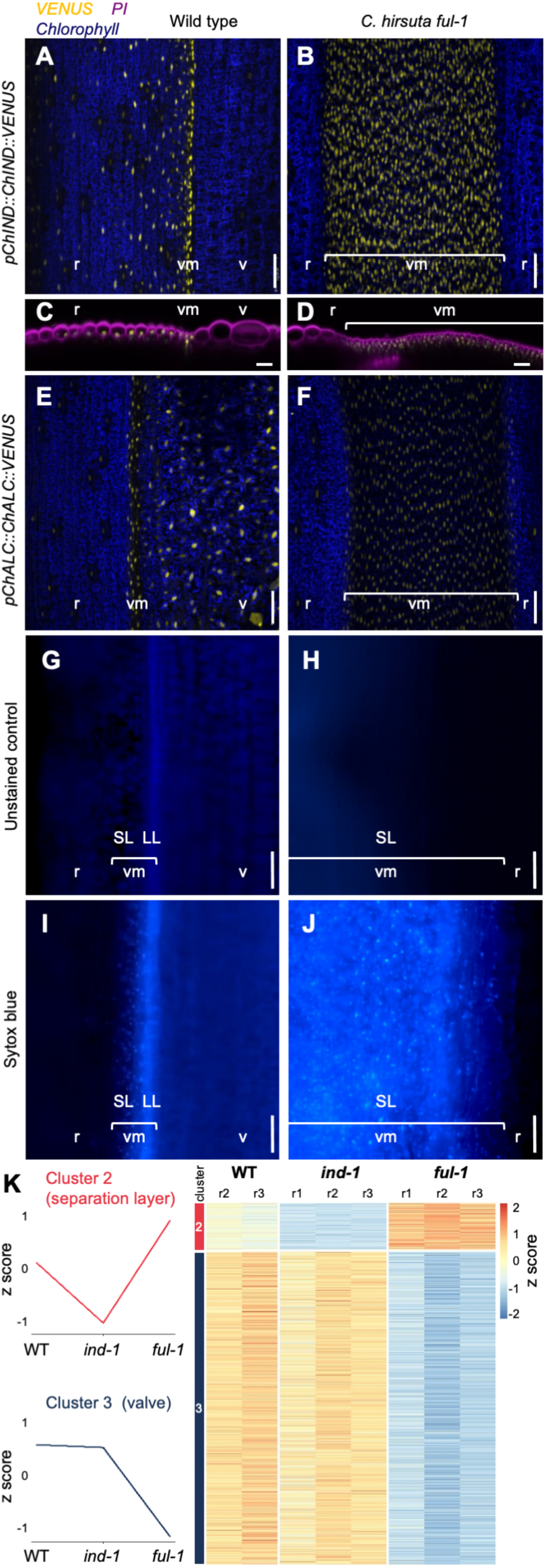
Homeotic conversion of valve to valve margin identity in *C. hirsuta ful* fruit. (A-F) CLSM of pChIND::cChIND:VENUS expression (yellow) (A-D) and pChALC::cChALC:VENUS expression (yellow) (E-F) in *C. hirsuta* stage 17a wild-type (A, C, E) and *ful-1* (B, D, F) fruit; chlorophyll autofluorescence (blue, A-B, E-F); optical transverse slices through propidium iodide-stained z-stacks (magenta, C-D). (G-J) Fluorescence micrographs of *C. hirsuta* stage 17a wild-type (G, I) and *ful-1* (H, J) fruit either unstained (G-H) or stained with Sytox blue cell death maker (I-J). (K) RNAseq analysis of *C. hirsuta* wild-type, *ind-1* and *ful-1* stage 17b fruit. Two gene expression clusters were selected and compared to differentially expressed genes between wild type and *ind-1* (cluster 2, 97 genes), or between wild type and *ful-1* (cluster 3, 1367 genes); absolute logFC >1 and adjusted p-value <0.05. Heatmap indicates relative expression of each gene in each cluster, in each replicate sample of wild type, *ind-1* and *ful-1*. Abbreviations: v, valve; r, replum; vm, valve margin; LL, lignified layer; SL, separation layer. Scale bars: 50μm (A-J).

**Figure 5.**
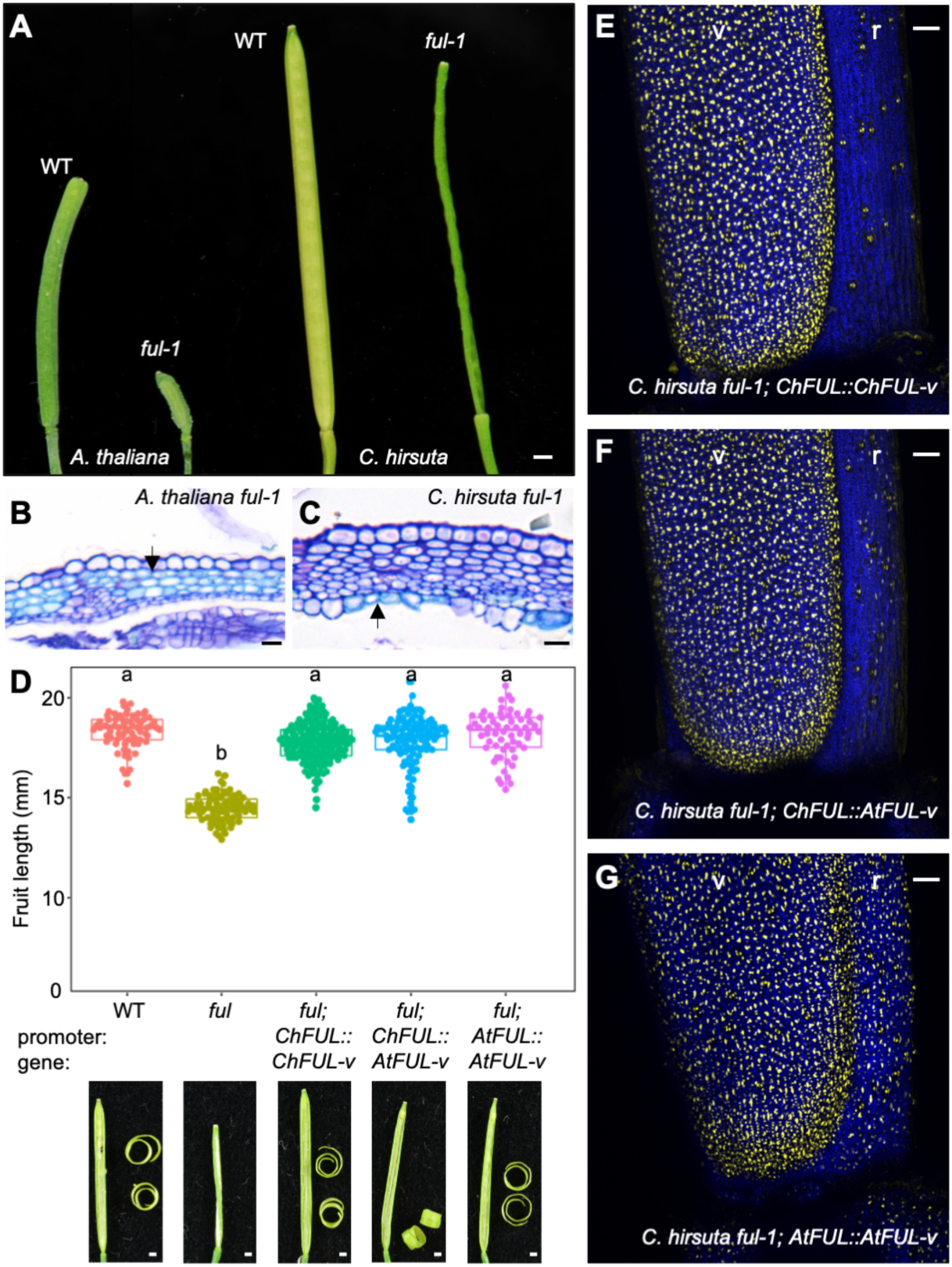
Conservation of *FUL* function between *A. thaliana* and *C. hirsuta*. (A) Mature fruit of *A. thaliana* Landsberg *erecta* wild type, *ful-1*, and *C. hirsuta* wild type*, ful-1*. (B-C) TBO-stained transverse sections of stage 17 *A. thaliana ful-1* (B) and *C. hirsuta ful-1* (C), arrows indicate lignified cell walls stained cyan. (D) Beeswarm boxplot of final fruit length and photographs of dehiscence in *C. hirsuta* wild type, *ful-1*, and *ful-1* complemented with *pChFUL::gChFUL-Venus*, *pChFUL::gAtFUL-Venus*, and *pAtFUL::gAtFUL-Venus*, coiled valves are shown with dehisced fruit. N = > 15 fruit from at least 5 plants for each genotype. Letters denote statistically significant differences (P < 0.001) between means based on one-way ANOVA (Tukey’s HSD). (E-G) CLSM showing FUL expression (yellow) in the fruit base of *C. hirsuta ful-1* complemented with *pChFUL::gChFUL-Venus* (E), *pChFUL::gAtFUL-Venus* (F), and *pAtFUL::gAtFUL-Venus* (G); chlorophyll autofluoresence in blue. Representative T2 lines of 10 independent transgenic lines shown in (E, F) and progeny of a representative T1 line crossed to *ful-1* shown in (G). Abbreviations: v: valve, r: replum. Scale bars: 1 mm (A, D) 10 μm (B, C), 50 μm (E-G).

### Homeotic conversion of valve to separation layer fate in *C. hirsuta ful* mutant

To investigate whether valve identity has switched to dehiscence zone identity in *C. hirsuta ful-1*, we examined the expression of valve margin identity genes in the mutant. *pChIND::ChIND:VENUS* accumulates in the valve margin but is strictly absent from outer layers of the valve in wild-type fruit (Fig. 4A, C, Fig. 2J-K). In *C. hirsuta ful-1*, *pChIND::ChIND:VENUS* accumulates throughout the ectopic valve margin (Fig. 4B, D), indicating a homeotic switch in cell fate from valve to valve margin. We used a *pAtIND::AtIND:GUS* reporter to show that ectopic *IND* expression is already obvious by stage 12 in *C. hirsuta ful-1* (Fig. S6), which coincides with the first differences in histology between *C. hirsuta ful-1* mutant and wild-type fruit (Fig. S6).

A second bHLH protein that acts in the valve margin in *A. thaliana* is ALC (Rajani and Sundaresan, 2001). We observed strong expression of *pChALC::ChALC:VENUS* in the valve margin of wild-type *C. hirsuta* fruit and throughout the ectopic valve margin in *C. hirsuta ful-1* (Fig. 4E-F). In wild-type fruit, we observed a sharp boundary between expression of *C. hirsuta* ALC in the valve margin and its absence from the replum, with expression extending from the valve margin part way into the valve (Fig. 4E). We also observed non-specific expression of *pChALC::ChALC:VENUS* in stomata throughout the fruit (Fig. 4E). A third gene that confers valve margin identity in *A. thaliana* is the MADS-box gene *SHP2* (Liljegren *et al*., 2000). We found expression of *pChSHP2::GUS* in the valve margin of wild-type fruit and throughout the ectopic valve margin in *C. hirsuta ful-1* (Fig. S6). Therefore, our results using three different marker genes confirm that valve cell fate is replaced by dehiscence zone fate in *C. hirsuta ful-1* fruit.

The dehiscence zone is comprised of two cell types: lignified and separation layer cells (Fig. 1). We hypothesized that the small, non-lignified cells found in the expanded dehiscence zones in *C. hirsuta ful-1* are separation layer cells (Fig. 3O). To test this, we used the Sytox Blue dead cell stain to identify separation layer cells, which are dead at maturity, in wild-type and *C. hirsuta ful-1* fruit. Under UV excitation, we observed bright speckles of Sytox Blue fluorescence in the separation layer of the wild-type dehiscence zone and throughout the expanded dehiscence zones in *C. hirsuta ful-1* (Fig. 4I-J). We also observed lignin autofluorescence in the lignified layer of the wild-type dehiscence zone in both the absence and presence of stain, but not in the expanded dehiscence zones of *C. hirsuta ful-1* (Fig. 4G-I). Therefore, the dehiscence zones that replace valve tissue in *C. hirsuta ful-1* comprise mostly separation layer rather than lignified layer cells.

To further characterize separation layer identity in *C. hirsuta* fruit, we generated transcriptional profiles of stage 17ab wild type, *ind-1*, and *ful-1* fruit. Seeds were dissected out of the fruit and not included. To identify valve margin genes that are likely to be associated with the separation layer, we combined differential expression and cluster analysis (Fig. 4K). Given that the ectopic valve margin tissue in *C. hirsuta ful-1* has separation layer identity, we predicted that genes downregulated in *ind-1*, but upregulated in *ful-1* may be associated with the separation layer. We selected cluster 2 from eight clusters identified in the data, because it showed this predicted pattern of gene expression (Fig. 4K, Fig. S7, Table S3). By comparing cluster 2 with 558 genes that are differentially expressed between wild type and *ind-1* (Table S4), we identified 97 shared genes (Fig. 4K, Table S5, Fig. S7). We used gene ontology (GO) analysis to determine that the most significantly enriched process in this gene set is transcriptional regulation (3 GO terms including GO:0006355, p-value 5.43E-04; Table S6). These GO terms include the related bHLH transcription factor genes *IND*, *SPT*, *HECATE2* and *HEC3* (Table S6). Cell wall organization is another process enriched in this gene set (GO:0071555, p-value 0.092), and includes the polygalacturonase gene *ADGP1* that is required to degrade pectins and allow cell separation along the dehiscence zone (Ogawa *et al*., 2009). Therefore, the separation layer transcriptome of stage 17ab *C. hirsuta* fruit is enriched for transcription factors and cell wall modifying enzymes.

We followed a similar approach to characterize valve tissue identity. By clustering genes that are similarly expressed between *C. hirsuta* wild type and *ind-1*, but downregulated in *ful-1*, we aimed to enrich for genes associated with the absence of valve, rather than the presence of ectopic valve margin in *C. hirsuta ful-1.* We selected cluster 3 because it showed this predicted pattern of gene expression (Fig. 4K, Fig. S7, Table S3). By comparing cluster 3 with 3505 genes that are differentially expressed between wild type and *ful-1* (Table S7), we identified 1367 genes in common (Fig. 4K, Table S8, Fig. S7). *FUL* is one of the most significantly differentially expressed genes in this cluster, and contributes, together with other transcription factor and receptor kinase genes, to the enriched GO term cell differentiation (GO:0030154, p-value 0.011, Table S9). The most significantly enriched processes in cluster 3 are associated with the cell wall (GO:0071555, p-value 9.13E-13), secondary cell wall (GO:0009834, p-value 3.56E-12), lignin biosynthesis (GO:0009809, p-value 1.1E-9), and cuticle development (GO:0042335, p-value 2.79E-4), suggesting that major cell wall remodelling occurs during the differentiation of valve-specific cell types (Table S9). Photosynthesis is another process enriched in this gene set (GO:0015979, p-value 1.95E-5), suggesting a high contribution of the valve to the photosynthetic activity found in fruit pods (Brazel and O’Maoileidigh, 2019). Therefore, the valve transcriptome of stage 17ab *C. hirsuta* fruit reflects the differentiation and function of valve-specific cell types.

### Conservation of *FUL* function between *A. thaliana* and *C. hirsuta*

In both *C. hirsuta* and *A. thaliana*, *FUL* functions in the fruit to specify valve identity by repressing valve margin identity genes such as *IND* (Fig. 4A-D) (Ferrandiz *et al*., 2000b; Liljegren *et al*., 2004). Yet *FUL* loss of function has different consequences for final fruit morphology in the two species (Fig. 5A). The strong *ful-1* allele in *A. thaliana* has severely stunted fruit valves with twisted repla (Gu *et al*., 1998), which differs dramatically from the straight, elongated fruit of *C. hirsuta ful-1* (Fig. 5A, Fig. S8). An allelic series in *A. thaliana* shows that *ful* phenotypes differ according to allelic severity in the extent to which the valves are replaced by dehiscence zone identity (Ferrandiz *et al*., 2000a). This suggests that the complete homeotic conversion of valve to dehiscence zone identity in the elongated fruit of *C. hirsuta ful-1* is likely to represent a strong, rather than weak, *ful* phenotype.

An important difference between the ectopic dehiscence zone that forms in *A. thaliana* versus *C. hirsuta ful-1* mutants is the cell type identity. The majority of this ectopic tissue has separation layer cell identity with some lignified cells in *C. hirsuta ful-1,* while in *A. thaliana ful-1,* this tissue has mostly lignified layer cell identity with some non-lignified, separation layer cells (Fig. 5B-C, Fig. S8). This suggests that although both species produce wild-type dehiscence zones with lignified and separation layers (Fig. 1), the ectopic dehiscence zone formed in the absence of *FUL*, is shifted towards separation layer identity in *C. hirsuta* and lignified layer identity in *A. thaliana*. To investigate whether this reflects a degree of divergence in *FUL* function between species, we complemented *ful-1* mutants from *C. hirsuta* and *A. thaliana* with the *FUL* genomic locus from both *C. hirsuta* and *A. thaliana*. We found that a *pChFUL::gChFUL:Venus* transgene, containing 6800 bp of upstream promoter sequence, complemented fruit length and dehiscence in *C. hirsuta ful-1* (Figs. 3C, 5D). We observed similar complementation of *C. hirsuta ful-1* with a *pAtFUL::gAtFUL:Venus* transgene containing 1893 bp of upstream promoter sequence (Fig. 5D). By generating chimeric constructs between the *FUL* promoter and gene sequences of each species, we showed that fruit of *C. hirsuta ful-1* were fully complemented by a *pChFUL::gAtFUL:Venus* transgene (Fig. 5D). Therefore, *C. hirsuta* and *A. thaliana FUL* genes are both sufficient to complement *C. hirsuta ful-1.* These experiments also demonstrated that the *FUL* promoter sequence caused a small but consistent difference in fruit length between *A. thaliana ful-1* complementation lines (Fig. S9). The 6800 bp promoter sequence of *C. hirsuta FUL* restored *A. thaliana ful-1* fruit length closer to wild type than the 1893 bp promoter sequence of *A. thaliana FUL* (Fig. S9), suggesting that the shorter promoter sequence is insufficient for full complementation. We found that the complementing FUL-Venus fusion proteins expressed strongly in the valves of *C. hirsuta* fruit (Fig. 5E-G). The 6800 bp promoter sequence of *C. hirsuta FUL* directed valve-specific expression of both *C. hirsuta* and *A. thaliana FUL*, in addition to non-specific expression in stomata (Fig. 5E-F). In comparison, expression driven by the shorter *A. thaliana FUL* promoter was not valve-specific, but extended through the valve margin and replum (Fig. 5G), suggesting that this 1893 bp promoter sequence is insufficient to drive correct expression. Overall, these results indicate a high degree of conservation in *FUL* function between *C. hirsuta* and *A. thaliana*.

### *C. hirsuta IND* is sufficient for separation and lignified layer cell types

To investigate whether the ectopic expression of *IND* in *C. hirsuta ful-1* is responsible for the ectopic dehiscence zones that replace valves in this mutant, we generated *ful-1; ind-1* double mutants in *C. hirsuta*. Double mutant fruit resembled *ind-1* single mutants; valve-like tissue with epidermal stomata, expanded cells and vascular bundles, replaced the ectopic dehiscence zones found in *C. hirsuta ful-1* (Fig. 6A-H, L-M, O). Therefore, ectopic *IND* expression is responsible for the ectopic dehiscence zones, which have mostly separation layer cell fate, in *C. hirsuta ful-1*.

**Figure 6.**
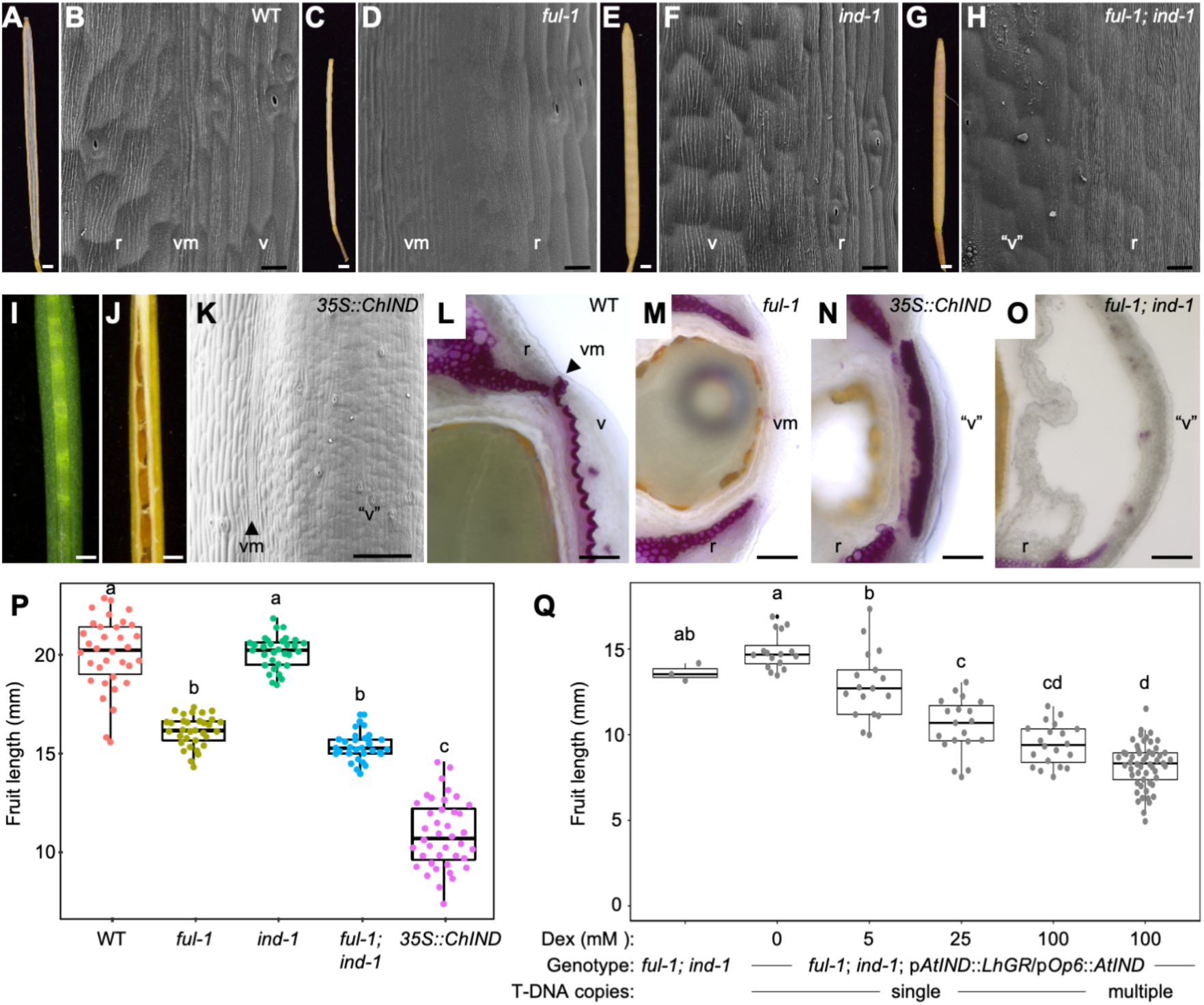
*C. hirsuta ful* phenotype is *IND*-dependent. **(A-K)** *C. hirsuta* fruit and SEMs of the fruit surface in dehiscent wild type (A, B) and indehiscent *ful-1* (C, D), *ind-1* (E, F), *ful-1;ind-1* (G, H), and *35S::ChIND* (I-K). (L-O) Phloroglucinol stained transverse sections of *C. hirsuta* fruit of the following genotypes: wild type (L), *ful-1* (M), *35S::ChIND* (N) and *ful-1;ind-1* (O). (P) Beeswarm boxplot of final fruit length in *C. hirsuta* wild type, *ful-1*, *ind-1*, *ful-1;ind-1*, and *35S::ChIND*, n = > 30 fruit from 5 plants for each genotype including 4 independent *35S::ChIND* T2 lines. (Q) Beeswarm boxplot of final fruit length in *C. hirsuta ful-1;ind-1* and *ful-1;ind-1; pAtIND::LhGR/pOp6::AtIND* homozygous T3 plants with single or multiple T-DNA insertions, treated with dexamethasone concentrations of 0, 5, 25 or 100 μM, n = 128 fruit from 4 (single T-DNA) and 8 (multiple T-DNA) independent transgenic lines. Letters denote statistically significant differences (P < 0.05) between means based on one-way ANOVA (Tukey’s HSD). Abbreviations: r, replum; v, valve; vm, valve margin; “v”. modified valve. Scale bars: 1 mm (A, C, E, G), 0.5 mm (I-J), 20 μm (B, D, F, H), 100 μm (K), 50 μm (L-O).

To investigate whether *C. hirsuta IND* is sufficient to specify the lignified layer cell type of dehiscence zones, we expressed *C. hirsuta IND* broadly in wild-type plants using the CaMV *35S* promoter. These *p35S::ChIND* fruit look like *C. hirsuta ful-1* fruit with non-explosive valves that resemble dehiscence zone tissue (Fig. 6I). But unlike *C. hirsuta ful-1*, the valves of *p35S::ChIND* fruit are ectopically lignified, indicating that *C. hirsuta IND* is sufficient to specify the lignified layer cell type of dehiscence zones (Fig. 6L-N). The lignified valves of *p35S::ChIND* fruit retain epidermal features of valve tissue, such as stomata (Fig. 6K). These fruits also form valve margins that lack stomata and comprise smooth, elongated, cells (Fig. 6K). The presence of non-lignified, separation layer cells at the valve margin may explain why *p35S::ChIND* fruit are not indehiscent like *C. hirsuta ful-1* but form a functional dehiscence zone between the replum and the lignified valves (Fig. 6J, N). Fruit length is significantly reduced in *p35S::ChIND* and is associated with ectopic lignification of the valve (Fig. 6N, P). This result lends support to the idea that the lignified layer cells that form in the ectopic dehiscence zones of *A. thaliana ful-1* fruit contribute to their severe reduction in fruit length (Fig. 5A-B, Fig. S8, Fig. S9) (Liljegren *et al*., 2004). However, *C. hirsuta ful-1* fruit also show a slight but significant reduction in length compared to wild type, which is not restored in *C. hirsuta ful-1; ind-1* double mutants (Fig. 6P). This difference in fruit length is not associated with significant differences in seed number between genotypes (Fig. S9), suggesting that it is not caused by differential seed set (Cox and Swain, 2006; Ripoll *et al*., 2019). Therefore, *FUL* may have a small but significant effect on fruit growth in *C. hirsuta* that is independent of *IND*, similar to *A. thaliana* (Ripoll *et al*., 2015).

To further investigate the link between ectopic *IND* gene expression and fruit length, we induced *IND*, driven by its own promoter, in a *ful-1; ind-1* background in *C. hirsuta.* Given that *A. thaliana IND* fully complements the *C. hirsuta ind-1* mutant (Fig. S10), we took advantage of a dexamethasone-inducible construct of *A. thaliana IND* (pAtIND:GR-pOp6:AtIND (Li *et al*., 2018)). Using increasing concentrations of dexamethasone to induce *IND* expression, we found a corresponding decrease in fruit length (Fig. 6Q). This quantitative relationship between *IND* expression and fruit length was associated with the formation of small valve margin cells in place of the larger valve-like cells observed in mock-treated fruit (Fig. S10). Moreover, the ectopic valve margin tissue in the shortest fruit, caused by high levels of *IND* expression, contained a greater proportion of lignified cells (Fig. S10). This indicates that different levels of ectopic *IND* expression are associated with different dehiscence zone cell fates. For example, we found in *C. hirsuta* that high levels of ectopic *IND* expression (*35S::ChIND*) causes mostly lignified layer cell fate, while much lower levels of ectopic *IND* (*C. hirsuta ful-1*) causes mostly separation layer cell fate, and intermediate levels of ectopic *IND* (dexamethasone-treated *C. hirsuta ful-1; ind-1; pAtIND:GR-pOp6:AtIND*) causes a mixture of both cell fates (Fig. 6, Fig. S10). These findings suggest that levels of IND activity may contribute to the specification of valve margin cells as having lignified or separation layer fates.

## Discussion

We showed that dehiscence zone formation is regulated by the bHLH protein IND in *C. hirsuta*, and is necessary for explosive fracture of the fruit. Without dehiscence zones, the fruit of *C. hirsuta ind* mutants are non-explosive. However, these mutant fruit valves coil when manually separated from the replum, indicating that they generate tension. Therefore, formation of a dehiscence zone along the valve margin is required to weaken the attachment between valve and replum in order to release this tension by valve coiling. Thus, by patterning dehiscence zone cell fate, IND regulates explosive seed dispersal in *C. hirsuta*.

Explosive seed dispersal is a key trait that distinguishes *C. hirsuta* from its close relative *A. thaliana*. Through a forward genetics screen for indehiscent mutants in *C. hirsuta*, we recovered *FUL* as a negative regulator of genes that promote dehiscence zone fate at the valve margins, including *IND*, *ALC* and *SHP2*. Therefore, the gene regulatory networks that pattern explosive versus non-explosive fruit tissues are conserved between *C. hirsuta* and *A. thaliana*, but we also identified divergence. Loss of *FUL* function results in cell fate transformations from valve to dehiscence zone in each species, but this fate is predominantly that of lignified layer cells in *A. thaliana,* and separation layer cells in *C. hirsuta.* This difference has profound consequences for *ful* fruit morphology between species, resulting in severely stunted fruit with twisted repla in *A. thaliana* (Gu *et al*., 1998), versus straight, elongated fruit in *C. hirsuta*. We show a quantitative relationship between ectopic *IND* expression and fruit length in *C. hirsuta* that correlates with the extent of ectopic lignification. Thus, the transformation to small, dehiscence zone cells with lignified layer fate impedes fruit expansion in both species (Liljegren *et al*., 2004). In this way, *ful* mutant fruit look very different between *C. hirsuta* and *A. thaliana* due to differences in the regulation of IND activity that are revealed in the absence of repression by FUL.

Overall conservation of the fruit patterning network between *C. hirsuta* and *A. thaliana* allows both species to produce equivalent patterning outputs despite divergence over evolutionary time – a concept called developmental system drift (Homann *et al*., 2009; True and Haag, 2001; Wotton *et al*., 2015). In order for system drift to occur, there must be different network genotypes that produce the same patterning outcome. For example, the repression of *IND* by FUL is conserved in *A. thaliana, C. hirsuta* and *Capsella rubella,* and is necessary for patterning the dehiscence zone in all three species (Dong *et al*., 2019; Liljegren *et al*., 2004). But FUL-independent regulation of *IND* may diverge more readily between species. For example, *cis-*regulatory changes in *Capsella rubella IND* cause its expression domain to expand into the fruit valves, where it activates the auxin biosynthesis genes *TAA1* and *YUCCA9* to influence fruit shape (Dong *et al*., 2019). A SUMO protease, HEARTBREAK, also participates in this process and stabilises *Capsella rubella* IND protein levels via de-SUMOylation (Dong *et al*., 2020). Therefore, both the levels and domain of IND expression can be regulated independently of FUL and diverge between species. In this study, we showed that manipulating *IND* expression levels shifts the balance of dehiscence zone cell fates. By quantitatively increasing the ectopic expression of *IND* in *C. hirsuta* fruit, we re-created the differences observed between elongated fruit lacking ectopic lignin, that are typical of *C. hirsuta ful*, and stunted fruit with ectopic lignin, which typify *A. thaliana ful*. This suggests that system drift between *C. hirsuta* and *A. thaliana* may have produced different network genotypes that quantitatively affect *IND* gene expression.

A key output of the fruit patterning network is to produce sharp borders between tissues. In *A. thaliana* fruit, IND expression reflects these sharp tissue boundaries, being restricted to the valve margin and excluded from both valve and replum (Li *et al*., 2018; Liljegren *et al*., 2004). However, in *C. hirsuta* fruit, the valve margin expression of *C. hirsuta* IND and ALC only produced a sharp boundary on one side of the valve margin. Specifically, *C. hirsuta* IND expression is excluded from the valve, but extends into the replum, while ALC expression is excluded from the replum, but extends into the valve (Fig. 4A, C, E). Therefore, the sharp borders of the dehiscence zone in *C. hirsuta* fruit are not precisely reflected by the expression domains of either IND or ALC alone. Full complementation of *C. hirsuta ind-1* by the more restricted valve margin expression of *A. thaliana* IND (Fig. S10), suggests that this is the functional domain of IND expression in *C. hirsuta* fruit. However, these expression differences could also reflect how the same patterning output is produced in different ways in *C. hirsuta* vs *A. thaliana* fruit. For example, the combination of both IND and ALC may be required to define a sharp border on both sides of the dehiscence zone in *C. hirsuta*.

We gained new insights into the transcriptomes of separation layer and valve tissues of the fruit by leveraging both *ind* and *ful* mutants in *C. hirsuta*. The separation layer transcriptome was characterized by transcription factors and cell wall enzymes, including genes with known functions in this cell type such as *IND*, *SPT* and *ADPG1* (Girin *et al*., 2011; Liljegren *et al*., 2004; Ogawa *et al*., 2009). Other genes in the enriched cell wall term GO:0071555, such as the xyloglucan endotransglucosylase/hydrolases *XTH3* and *XTH28* (Table S6), could be investigated for possible functions in the separation layer or as markers for this cell type in *C. hirsuta* fruit. Similarly, transcripts such as *FUL* that were enriched in the valve, reflected the differentiation and function of valve-specific cell types. Future studies could investigate genes in the enriched lignin biosynthesis term GO:0009809, such as *LACCASE4, 11* and *17* (Table S9), as possible markers for the asymmetrically lignified endocarp *b* cell type in *C. hirsuta* fruit valves. Approximately 10% of the top 50 differentially expressed genes in each cluster lacked an annotation or a clear ortholog in *A. thaliana* (Tables S5, S8), which may indicate a role for *C. hirsuta*-specific genes in fruit valve and separation layer development. Identifying new genes that act in the explosive fruit of *C. hirsuta*, and how these genes act in relation to *IND* and *FUL*, will further our understanding of how the balance of conservation versus divergence in fruit gene regulatory networks produced different seed dispersal strategies during evolution.

## Materials and Methods

### Plant material and growth conditions

*C. hirsuta* reference Oxford (Ox) accession, herbarium specimen voucher Hay 1 (OXF) (Hay and Tsiantis, 2006), and *A. thaliana* Landsberg *erecta* accession were used throughout unless indicated. Gene identifiers in the *C. hirsuta* genome assembly (Gan et al., 2016): *FUL*, CARHR274590; *IND*, CARHR248170; *ALC,* CARHR280540; *CLATHRIN/AP2M*, CARHR174880; *PHYTOENE DESATURASE*, CARHR220290. *A. thaliana ful-1* mutant (Gu et al., 1998) was obtained from the Arabidopsis Biological Resource Center, accession number CS3759. All plants were grown in long day conditions in the greenhouse: 16 h at 22°C: 8 h at 20°C, light: dark.

### EMS mutagenesis, mutant screening, genetic mapping and gene identification

*C. hirsuta* seeds were washed in 0.1% Triton® X-100, treated with either 0.18% (15 mM) or 0.36% (30 mM) ethyl methane sulphonate (EMS) in water for 10 hours, repeatedly washed with water, and pipetted onto soil as previously described (Hofhuis et al., 2016). Seed was collected from all fertile M1s, giving 895 M2 lines, of which 32145 M2 seeds were planted and screened for indehiscence. To map the indehiscent *ful-1* mutation, homozygous mutants were crossed to the Wa accession and F1 plants were selfed to generate 743 F2 plants. DNA was extracted from 148 *ful-1* mutants in this F2 and the mutant locus was mapped to a 11.7 cM interval flanked by PCR markers m229 and m306 at the bottom of chromosome 8. Further mapping delimited a 2.7 cM interval flanked by PCR markers fd04 and m690, which corresponded to a region of chromosome 5 in *A. thaliana* that contained 33 annotated genes (TAIR10), including *FUL,* AT5G60910. Sanger sequencing identified a C to T single nucleotide change at position 429 of the full length *FUL* CDS in *C. hirsuta ful-1*, predicted to convert a Gln residue to a stop codon and produce a truncated 143 amino acid protein. A CAPS marker was designed to distinguish this SNP from the wild type Ox and Wa sequence and was shown to co-segregate with the *ful-1* phenotype in *C. hirsuta*. **CRISPR-Cas9 mutagenesis.** SgRNA-*IND* (5’-AAGCGACGATCCTCAGACGGTGG-3’), located upstream of the HEC domain in *C. hirsuta IND*, was subcloned into pYB196 (Hyun et al., 2015) to generate a pU6:sgRNA-ChIND/pICU2:Cas9 T-DNA cassette. Floral dip was used to transfer this cassette into *C. hirsuta* by *Agrobacterium tumefaciens*-mediated transformation. T1 transformants were selected as Basta-resistant, the sgRNA flanking region was PCR-amplified, Sanger-sequenced and analyzed for induced mutations using http://yosttools.genetics.utah.edu/PolyPeakParser to parse double peaks into wild-type and mutant allele sequences. A single T-DNA insertion line contained a nucleotide insertion at position 343 bp from the first Met that converted the GAC triplet into GAAC, introducing a frameshift. T2 progeny of this plant were PCR genotyped with M13F and sg1 primers to identify and discard transgenic plants, then the transgene-free plants were screened for the expected indehiscent fruit phenotype. We confirmed that indehiscent plants were homozygous for the nucleotide insertion described above using Ta-sensitive PCR primers (chind-wt-F, chind-mutant–F and chind-qRT-R) and HiDi DNA polymerase. The allele was named *C. hirsuta ind-1*.

### Construction of transgenes

pAtIND::AtIND:EYFP and pAtIND:GR-pOp6:AtIND were a kind gift from L. Ostergaard (Li et al., 2018) and gAtFUL:GFP from G. Angenent (Urbanus et al., 2009). MultiSite Gateway cloning was used to generate all other constructs unless indicated. pChIND::ChIND:Venus: 1,885 bp of untranscribed *IND* promoter sequence, corresponding to the entire intergenic region, was amplified from genomic *C. hirsuta* DNA with PCR primers pChIND-attB4-F and pChIND-attB1r-R, and subcloned into pGEM1R4_easy entry vector; 585 bp of transcribed *IND* sequence from the first Met and excluding the STOP codon was amplified from genomic *C. hirsuta* DNA with PCR primers ChIND-attB1-F and ChIND-attB2-R, and subcloned into pGEM221_easy entry vector; these entry vectors were recombined with the p4gly-vYFP-3AT_2R3 entry vector to generate pChIND::ChIND:Venus in the pGREENII125 binary vector that contains norflurazon selection (Galinha et al., 2007; Prasad et al., 2011). pChIND::3xVenus: pChIND_1R4 and 3x-vYFP_221 entry vectors were recombined to generate pChIND::3xVenus in the pGREENII125 binary vector. pChIND-UTR::3xVenus: 1,969 bp sequence including 1,885 bp *IND* promoter and an additional 84 bp of predicted 5’UTR that corresponds to the first 28 amino acids shown in Figure S2B, was amplified from genomic *C. hirsuta* DNA with PCR primers pChIND-attB4-F and pChIND-attB1r-R-UTR, subcloned into pGEM1R4_easy entry vector, and recombined with 3x-vYFP_221 entry vector to generate pChIND-UTR::3xVenus in the pFANTASTIC (pFAN) binary vector that contains a pOLE1:OLE1:RFP cassette for seed fluorescence selection. pFAN is a modified version of pPZP200 containing the R4-R3 MultiSite Gateway cassette from pGREENII125. p35S::ChIND: 585 bp of transcribed *IND* sequence from the first Met and including the STOP codon was amplified from genomic *C. hirsuta* DNA with PCR primers (ChIND-F-XhoI and ChIND-R-BamHI), subcloned into pART7 using *<u>Xho</u>*I and *Bam*HI restriction enzymes to generate p35S:ChIND-NosT, and transferred as a *Nco*I fragment into the binary vector pGREENII125

pChFUL::ChFUL:Venus: 6800 bp of *FUL* promoter sequence was amplified from genomic *C. hirsuta* DNA with PCR primers pChFUL-attB4-F and pChFUL-attB1r-R, and subcloned into pGEM1R4_easy entry vector; 3400 bp of *FUL* gene including 5’UTR but excluding the STOP codon was amplified from genomic *C. hirsuta* DNA with PCR primers gAtFUL-attB1-F and gChFUL-attB1-R, and subcloned into pGEM221_easy entry vector; these entry vectors were recombined with the p4gly-vYFP-3AT_2R3 entry vector to generate pChFUL::ChFUL:Venus in pFAN. pChFUL::Gus: pChFUL_1R4 and GUS_221 entry vectors were recombined to generate pChFUL::Gus in the pGREENII125 binary vector. pAtFUL::gAtFUL:Venus: 1893 bp of *FUL* promoter sequence was amplified from gFUL:GFP (de Folter et al., 2007) with PCR primers pAtFUL-attB4-F and pAtFUL-attB1r-R, and subcloned into pGEM1R4_easy entry vector; 3400 bp of *FUL* gene including 5’UTR but excluding the STOP codon was amplified from genomic *A. thaliana* L.*er* DNA with PCR primers gAtFUL-attB1-F and gAtFUL-attB2-R, and subcloned into pGEM221_easy entry vector; these entry vectors were recombined with the p4gly-vYFP-3AT_2R3 entry vector to generate pAtFUL::AtFUL:Venus in pFAN and pGREENII125 binary vectors. pChFUL::gAtFUL:Venus: pChFUL_1R4 and AtFUL_221 entry vectors were recombined with p4gly-vYFP-3AT_2R3 to generate pChFUL::AtFUL:Venus in pFAN and pGREENII125 binary vectors. pAtFUL::gChFUL:Venus: pAtFUL_1R4 and ChFUL_221 entry vectors were recombined with p4gly-vYFP-3AT_2R3 to generate pAtFUL::ChFUL:Venus in pFAN and pGREENII125 binary vectors. pChALC::ChALC:Venus: 720 bp of untranscribed *ALC* promoter sequence, corresponding to the entire intergenic region, was amplified from genomic *C. hirsuta* DNA with PCR primers AG119-pChALC-attB4-F and AG120-pChALC-attB1r-R, and subcloned into pGEM1R4_easy entry vector; 633 bp of transcribed *ALC* sequence from the first Met and excluding the STOP codon was amplified from genomic *C. hirsuta* DNA with PCR primers (AG129-ChALC-attB1-F and AG130-ChALC-attB2-R) and subcloned into pGEM221_easy entry vector (cChALC_221); these entry vectors were recombined with the p4gly-vYFP-3AT_2R3 entry vector to generate pChALC::ChALC:Venus in the pFAN binary vector that contains seed-coat-RFP fluorescence selection marker.

All plasmids are listed in Table S2. Floral dip was used to transfer plasmids into *C. hirsuta* and *A. thaliana* genotypes by *Agrobacterium tumefaciens*-mediated transformation. Transgene copy number was determined by g-Count (IDna Genetics Ltd) and 5-15 single copy lines were analyzed for each transgene to determine consensus phenotypes and expression patterns for each transgene.

### Virus Induced Gene Silencing (VIGS)

pTRV2-ChFUL-ChPDS was constructed by cloning a 421 bp fragment containing part of the K domain, the C domain, and part of the 3’UTR of *C. hirsuta FUL,* that was amplified from fruit cDNA using primers ChFUL-F and ChFUL-R, and cloned into pTRV2-ChPDS via EcoRI and XbaI restriction sites. pTRV2-ChPDS was constructed by cloning a 471 bp fragment of *C. hirsuta PHYTOENE DESATURASE* ChPDS (CARHR220290.1), that was amplified using primers ChPDS-F and ChPDS-R, into pTRV2 via XbaI and KpnI restriction sites. Both pTRV2-ChFUL-ChPDS and pTRV2-ChPDS were transformed into *Agrobacterium tumefaciens* GV3101 and VIGS was performed as previously described (Kramer et al., 2007) by infiltrating young *C. hirsuta* leaves.

### Histology

Plastic embedding and Toluidine blue O (TBO) staining of fruit tissue was similar to (Neumann and Hay, 2019). Fruit were fixed with 1% gluturaldehyde (Agar Scientific) and 4% formaldehyde (Fisher Scientific) in 0.05 M phosphate buffer (pH 7.2) for 15 minutes and then left shaking at 4°C overnight. After washing with 0.5 M phosphate buffer (pH 7.2), samples were dehydrated through an ethanol series and gradually embedded in Technovit 7100 resin at room temperature. The resin was polymerized and embedded with hardener I and hardener II according to manufacturer’s instructions and epoxy glued to rubber stubs. Semi-thin sections (5 μm) were collected on glass slides, stained with 0.05% TBO in 0.1 M phosphate buffer (pH 5.7), rinsed with dH_2_O and mounted in DPX Mounting Medium (Agar Scientific). Images were taken with a Zeiss Axio Imager.

Propidium iodide (5 mg/mL) staining was performed with either whole fruit segments for 10 min, or with thick vibratome-cut sections (150 µm) for 5 min, after fixing in 2% paraformaldehyde and embedding in 10% low melt agarose. Samples were immediately imaged with a Leica SP8 upright confocal microscope.

Staining with the dead cell marker Sytox Blue (S34857, Thermo Fisher) was performed by incubating fruit in 1 µM aqueous Sytox Blue in Eppendorf tubes for 6h at room temperature in the dark. Fruits were imaged in water using a Zeiss Axio Imager with a DAPI filter. Nuclear sytox blue stain could be distinguished from cell wall lignin autofluorescence.

For phloroglucinol staining, fruit were fixed in 2% paraformaldehyde, embedded in 10% low melt agarose, cut into thick vibratome sections (150 µm) and stained with 2% phloroglucinol in 95% ethanol for 10 min at room temperature, then acidified by transferring to 10 M HCl for 1 min, mounted and imaged immediately in 1 M HCl using a Zeiss Axio Imager with differential interference contrast.

β-Glucuronidase (GUS) staining of whole fruit was performed as previously described (Lilley et al., 2012). After staining at 37°C overnight, fruit were fixed with either FAA (50% ethanol, 3.7% formaldehyde, and 5% acetic acid) or 2.5% gluturaldehyde in phosphate buffered saline for 60 min. After repeated washing with phosphate buffered saline, samples were dehydrated through an ethanol series into histoclear, embedded in Paraplast Plus, and 8-10 µm sections were deparaffinized and imaged using a Zeiss Axio Imager with differential interference contrast.

### Dexamethasone treatments

Dexamethasone (Sigma-Aldrich) solutions of 5 µM, 25 µM and 100 µM were freshly prepared from a 10 mM aqueous stock with 0.015% Silwet L-77. Mock solutions contained 0.015% Silwet L-77 in water. Solutions were sprayed directly on *C. hirsuta* inflorescences just after bolting every 2 days over 2 weeks until the oldest fruit reached stage 17b.

### Scanning Electron Microscopy (SEM)

Fruit were fixed, dehydrated, critical point dried, sputter coated, and imaged with either a Zeiss Supra 40VP or a JSM-5510 (Joel) microscope as previously described (Neumann and Hay, 2019).

### Confocal laser scanning microscopy (CLSM)

A Leica SP8 upright CLSM with 20x and 40x objectives was used with these excitation wavelengths: Venus and EYFP (514 nm with 519-569 nm bandpass filter), tdTomato (554 nm with 600-680 nm bandpass filter), PI (488 nm with 628-735 nm bandpass filter), chlorophyll autofluorescence (514 with 628-735 nm bandpass filter).

### Time-lapse CLSM and quantitative image analysis

Time-lapse imaging was performed with *C. hirsuta pUBQ10::acyl:tdT* transgenic plants (gift from Tsiantis lab, modified from (Segonzac et al., 2012)). A reference point was gently marked in the centre of each fruit with permanent marker before and after imaging. Fruit were mounted in water with the valve margin facing the cover slip and imaged at 0 h (stage 15), 48 h (stage 16), 96 h (stage 17a) and 168 h (stage 17b). Fruit stages were estimated by fruit length: stage 15 = 5-9mm, stage 16 = 10-16 mm, stage 17 = 17-20 mm. Fruit remained attached to the plant, and plants were returned to the greenhouse between imaging. Image stacks were acquired at 512×512 pixels without averaging, with 0.5-0.8 μm distance in Z-dimension, and were analyzed using MorphoGraphX software (Barbier de Reuille et al., 2015). Multiple image stacks were required to capture later time points and these stacks were stitched using MorphoGraphX. After cells were segmented, parent relations between successive stages were determined manually and used to quantify cell area extension. Heat maps are displayed on the second of two time points. Cells were assigned tissue identities using the cell distance process with filtering based on cell morphology, and these identities were exported as parent labels to calculate area extension in organ-aligned directions in each tissue (Zhang et al., 2020).Two replicate time-lapse series were acquired from fruit on different plants in a single experiment, and progression through developmental stages to explosion was compared with non-imaged fruit to show a negligible effect of imaging on fruit development.

### Fruit measurements

Digital photographs of fruit (Nikon camera) were measured from stigma tip to gynophore base with the straight-line tool in FiJi (Schindelin et al., 2012).

### Quantitative RT-PCR analysis

*C. hirsuta* tissue samples were collected and immediately frozen in liquid nitrogen and stored at −80°C before processing. Total RNA was extracted using the Spectrum Plant Total RNA Kit (Sigma-Aldrich), treated with DNaseI on columns (Sigma-Aldrich), and converted into cDNA using SuperScript VILO (Thermo Fisher Scientific) and an oligo-dT primer. Quantitative PCR was performed in triplicate using Power SYBR Green Master Mix (Thermo Fisher Scientific) in a QuantStudio Real-Time PCR System (Applied Biosystems). Primer efficiency and gene expression were quantified using the standard curve method as previously described (Pfaffl, 2001). Primers were designed using the QuantPrime program and checked for specificity by BLAST to the *C. hirsuta* genome. Gene expression was normalized against the reference gene *CLATHRIN/AP2M* and estimated as fold change relative to a control sample, with error calculated using ΔCt values of biological replicates.

### RNA-seq analysis

Fruits of *C. hirsuta* wild type, *ind-1* and *ful-1* were harvested at stage 17b and seeds and septum tissue were removed. Total RNA was extracted from 100 mg of whole seedling tissue using the Spectrum Plant Total RNA Kit (Sigma, STRN50) and l μl of eluted RNA was used for complementary DNA (cDNA) synthesis using SuperScript Vilo (Invitrogene, 11754-050). RNA quality was assessed by Bioanalyser Nanochip (Agilent, Santa Clara, U.S.A.), and libraries quantified by Qubit (ThermoFisher, Waltham, U.S.A.). Libraries were prepared according to NEBNext Ultra™ Directional RNA Library Prep Kit for Illumina (New England Biolabs, Ipswich, U.S.A.) followed by single end sequencing on a HiSeq3000 to produce 2-3 Gb data per sample. RNA-sequencing reads were aligned against the *C. hirsuta* genome (Gan *et al*., 2016) and a cut-off of five reads in every sample that uniquely mapped to annotated genes was used as criteria for further analysis, giving a total of 22981 genes. We used the DESeq2 Bioconductor package (Anders and Huber, 2010) for all following analyses. Based on a principal component analysis of the log-transform of all samples, one wild-type replicate was removed (Fig. S7). We fitted a linear model to the genes to estimate the effect of each genotype (wild-type*, ind-1, ful-1*) on each gene. We then clustered the normalized, three-dimensional vectors into 8 different clusters (Fig. S7, Table S3). The number of clusters was selected based on the total variance within the clusters. Differentially expressed genes were identified between wild type and *ind-1*, and wild type and *ful-1* (Tables S4, S7). Then we applied a contrast analysis to obtain the differentially expressed genes in selected clusters (Tables S5, S8). We included genes with an absolute logFC higher than 1 and adjusted p-value lower than 0.05. *A. thaliana* othologs were retrieved for all *C. hirsuta* gene identifiers (Gan *et al*., 2016) and used with DAVID (Huang *et al* Nature 2009) for gene ontology analysis.

### Primers

All primer sequences are listed in supplementary Table S1.

## Acknowledgements

We thank A. Emonet for comments, W. Faigl, R. Berndtgen, S. Strauss, R. Lympouridou and S. Pophaly for valuable assistance, and L. Ostergaard and G. Angenent for materials.

## Author contributions

A.G. and A.H. designed experiments; A.G., P.S., L.N., M.P-A. and H.H. performed research; A.G., P.S., R.C-M., X.G., L.R-L. and M.A. analyzed data; A.H. designed and directed the study and wrote the paper with A.G.; all authors commented on the article.

## Data availability

ENA sequence read depository.

