## Supplemental Figures S1-S10 and Tables S1-S2 for "INDEHISCENT regulates explosive seed dispersal"

### Supplementary figures

#### Table S1 Primer list

| Purpose | Name | Identifier | 5' Sequence 3' | Method/Restriction enzyme | Description |
|---|---|---|---|---|---|
| ChIND-CRISPR | pU6-F-seq | AG69 | AGAATGATTAGGCATCGAAC | PCR amplification | Clone pU6:sgRNA-ChIND/pICU2:Cas9 |
| | sg-1 | AG70 | ATATACTAGTCTAGAGAATGATTAGGCATCGAACCT | PCR amplification | Clone pU6:sgRNA-ChIND/pICU2:Cas9 |
| | sg-2 | AG71 | ATGCAGGAAGACAACTAGTCAATAATC | PCR amplification | Clone pU6:sgRNA-ChIND/pICU2:Cas9 |
| | CRISPR-CHIND-F | AG72 | AAGCGACGATCCTCAGACGGGTTTTAGAGCTAGAAATA | PCR amplification | Clone pU6:sgRNA-ChIND/pICU2:Cas9 |
| | CRISPR-CHIND-R | AG73 | CCGTCTGAGGATCGTCGCTTCACAATCACTACTTCGACTCTA | PCR amplification | Clone pU6:sgRNA-ChIND/pICU2:Cas9 |
| genotype | ChIND-wt | AG141 | ATAAGCGACGATCCTCAGAC | HiDi polymerase PCR | Genotype *C. hirsuta ind-1* allele |
| | ChIND-mutant | AG142 | ATAAGCGACGATCCTCAGAA | HiDi polymerase PCR | Genotype *C. hirsuta ind-1* allele |
| genotype | cful4F | | GCTTCAAAGCTTGGAGCATC | DdeI | Genotype *C. hirsuta ful-1* allele |
| | cful4R | | GCAAGGCCTTATCCTGTTTG | DdeI | Genotype *C. hirsuta ful-1* allele |
| Mapping cful-1 | m229F | | TCCTTAGTTCGATATTTGCAAATC | Taqαl | CAPS marker |
| | m229R | | GATTTGAGAAGGAACCGTTGA | Taqαl | CAPS marker |
| | m306F | | TCCTTTGGGTCATCCTCTTG | DdeI | CAPS marker |
| | m306R | | CTTTCTCTTGCCGTGATGCT | DdeI | CAPS marker |
| | fd04F | | TATCGTGGTGTGCCGAGAACTG | PstI | CAPS marker |
| | fd04R | | AGCTTGACAACCCCATTCAC | PstI | CAPS marker |
| VIGS | ChPDS-F | | AAGATGGCATTCTTGGATGG | PCR amplification | Clone pTRV2-ChFUL-ChPDS/ pTRV2-ChPDS |
| | ChPDS-R | | GTCGCTCCGTGATATCCATT | PCR amplification | Clone pTRV2-ChFUL-ChPDS/ pTRV2-ChPDS |
| | ChFUL-F | | GGAATTTCATGGGGGAAGAT | PCR amplification | Clone pTRV2-ChFUL-ChPDS |
| | ChFUL-R | | GATGGATCGAGAGAGAGGTAATTC | PCR amplification | Clone pTRV2-ChFUL-ChPDS |
| qPCR | Clathrin-F | AG61 | TCGATTGCTTGGTTTGGAAGATAAGA | SyberGreen qPCR | house keeping gene for normalization |
| | Clathrin-R | AG62 | TTCTCTCCCATTGTTGAGATCAACTC | SyberGreen qPCR | house keeping gene for normalization |
| | qPCR-ChIND-F | AG77 | TTGGAGCTCCTATGGCTGACCC | SyberGreen qPCR | ChIND and AtIND |
| | qPCR-ChIND-R | AG78 | AAATCAGGCTTGGGAGTTGGGG | SyberGreen qPCR | ChIND and AtIND |
| | FUL-qRT-F2 | AG59 | TCGAATATTCCACCGACTCTTGC | SyberGreen qPCR | ChFUL |
| | FUL-qRT-R2 | AG60 | TTTGTGAAACGTCTCGGCCAAC | SyberGreen qPCR | ChFUL |
| Cloning | pChFUL-attB4-F | | GGGGACAACTTTGTATAGAAAAGTTGCTCAAAGCGAGAAGTTGCTT | Multisite gateway cloning | pChFUL_1R4 |
| | pChFUL-attB1r-R | | GGGGACTGCTTTTTTGTACAAACTTGCTCTTTTTTTCTTTGCTTTT | Multisite gateway cloning | pChFUL_1R4 |
| | gChFUL-attB1-R | AG08 | GGGGACCACTTTGTACAAGAAAGCTGGGTACTCATTAGTAGTAGGGCGT | Multisite gateway cloning | gChFUL_221, without STOP codon |
| | pAtFUL-attB4-F | AG01 | GGGGACAACTTTGTATAGAAAAGTTGCTCTTGTCGATCAGAATTTG | Multisite gateway cloning | pAtFUL_1R4 |
| | pAtFUL-attB1r-R | AG02 | GGGGACTGCTTTTTTGTACAAACTTGCATCTCTCTCTCTTCAAAAT | Multisite gateway cloning | pAtFUL_1R4 |
| | gAtFUL-attB1-F | AG03 | GGGGACAAGTTTGTACAAAAAAGCAGGCTTAATGGGAAGAGGTAGGGTT | Multisite gateway cloning | gAtFUL_221/ gChFUL_221 |
| | gAtFUL-attB2-R | AG04 | GGGGACCACTTTGTACAAGAAAGCTGGGTACTCGTTCGTAGTGGTAGGA | Multisite gateway cloning | gAtFUL_221, without STOP codon |
| | pChIND-attB4-F | AG127 | GGGGACAACTTTGTATAGAAAAGTTGCTGTCTAGAACTCATTAAGC | Multisite gateway cloning | pChIND_1R4 |
| | pChIND-attB1r-R | AG128 | GGGGACTGCTTTTTTGTACAAACTTGCCTTTTCTTATTTCTTAATA | Multisite gateway cloning | pChIND_1R4 |
| | ChIND-attB1-F | AG123 | GGGGACAAGTTTGTACAAAAAAGCAGGCTTAATGAAAACGGAAATTGGT | Multisite gateway cloning | cChIND_221 |

| | | | | |
|---|---|---|---|---|
| ChIND-attB2-R | AG124 | GGGGACCACTTTGTACAAGAAAGCTGGGTAGGCTTGGGAGTTGGGGTAA | Multisite gateway cloning | cChIND_221, without STOP codon |
| pChIND-attB1r-R-UTR | AG149 | GGGGACTGCTTTTTTGTACAAACTTGCGCTTCTTTTGGGGCTGTGG | Multisite gateway cloning | pChIND+UTR_1R4, with the 5' UTR, |
| ChIND-F-XhoI | AG57 | GGCTCGAGATGAAAACGGAAATTG | XhoI | cChIND |
| ChIND-R-BamHI | AG58 | CCGGATCCTCAGGCTTGGGAG | BamHI | cChIND with STOP codon |
| pChALC-attB4-F | AG119 | GGGGACAACTTTGTATAGAAAAGTTGCTGAGACCTCAAGTTGGATT | Multisite gateway cloning | pCALC_1R4 |
| pChALC-attB1r-R | AG120 | GGGGACTGCTTTTTTGTACAAACTTGCCTCTCTCTGTCTCTGTGTT | Multisite gateway cloning | pCALC_1R4 |
| chALC-attB1-F | AG129 | GGGGACAAGTTTGTACAAAAAAGCAGGCTTAATGGGTGATTCCGACGAC | Multisite gateway cloning | cChALC_221 |
| chALC-attB2-R | AG130 | GGGGACCACTTTGTACAAGAAAGCTGGGTAAAGCAGAGTGGGTGTGGGA | Multisite gateway cloning | cChALC_221, without STOP codon |

### Table S2 Plasmid list

| Name | Promoter | Gene | Vector | Reference | Species | Genotype |
|---|---|---|---|---|---|---|
| pU6:sgRNA-ChIND/pICU2: Cas9 | pICU2 | sgRNA-ChIND | pYB196 | this study | *C. hirsuta* | Ox |
| pChIND::ChIND:Venus | ChIND | ChIND:Venus | pGRENII125 | this study | *C. hirsuta* | *ind-1* |
| pChIND::3xVenus | ChIND | 3xVenus | pGRENII125 | this study | *C. hirsuta* | Ox |
| pChIND-UTR::3xVenus | ChIND-UTR | 3xVenus | pFANTASTIC | this study | *C. hirsuta* | Ox |
| pAtIND::AtIND:EYFP | AtIND | AtIND:EYFP | | Li XR *et al*. 2018. Molecular Plant | *C. hirsuta* | *ind-1* |
| pAtIND:GR-pOp6:AtIND | AtIND | GR/pOp6:AtIND | | Li XR *et al*. 2018. Molecular Plant | *C. hirsuta* | *ful-1;ind-1* |
| p35S::ChIND | 35S | ChIND | pGRENII125 | this study | *C. hirsuta* | Ox |
| pChFUL::Gus | ChFUL | Gus | pGRENII125 | this study | *C. hirsuta* | Ox |
| pChFUL::ChFUL:Venus | ChFUL | ChFUL:Venus | pFANTASTIC | this study | *C. hirsuta* | *ful-1* |
| pAtFUL::AtFUL:Venus | AtFUL | AtFUL:Venus | pFANTASTIC | this study | *C. hirsuta* | *ful-1* |
| pAtFUL::AtFUL:Venus | AtFUL | AtFUL:Venus | pGRENII125 | this study | *A.thaliana* | *ful-1* |
| pChFUL::AtFUL:Venus | ChFUL | AtFUL:Venus | pFANTASTIC | this study | *C. hirsuta* | *ful-1* |
| pChFUL::AtFUL:Venus | ChFUL | AtFUL:Venus | pGRENII125 | this study | *A.thaliana* | *ful-1* |
| pAtFUL::ChFUL:Venus | AtFUL | ChFUL:Venus | pFANTASTIC | this study | *C. hirsuta* | *ful-1* |
| pAtFUL::ChFUL:Venus | AtFUL | ChFUL:Venus | pGRENII125 | this study | *A.thaliana* | *ful-1* |
| pTRV2-ChPDS | TRV2 | ChPDS | pTRV2 | this study | *Agrobacterium tumifaciens* | |
| pTRV2-ChFUL | TRV2 | ChFUL | pTRV3 | this study | *Agrobacterium tumifaciens* | |
| pChALC::ChALC:Venus | ChALC | ChALC:Venus | pFANTASTIC | this study | *C. hirsuta* | Ox |

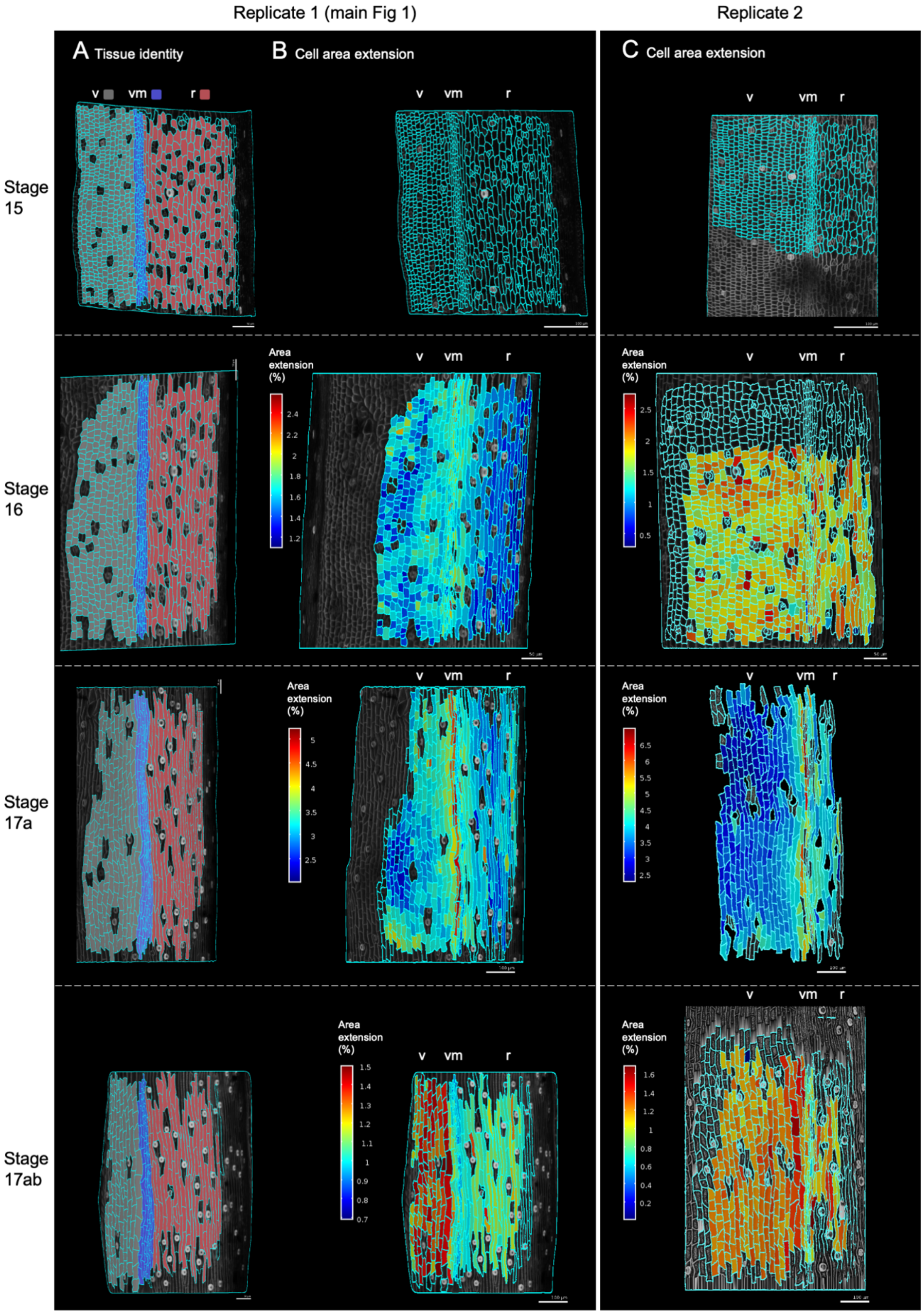

Replicate 1 (main Fig 1)
Replicate 2
A Tissue identity
B Cell area extension
C Cell area extension
v vm r
Stage 15
Stage 16
Stage 17a
Stage 17ab
Area extension (%)

**Fig. S1 Quantitative analysis of two time-lapse series of *C. hirsuta* fruit development from stage 15 to 17ab.** (A-B) Replicate 1 time-lapse series (also shown in Fig. 1P, Q main text). (A) Cells labelled by tissue identity (used for quantifications in main Fig. 1Q): valve (grey), valve margin (blue), replum (red). (B) Cell area extension (%) between consecutive stages, mapped on second time point (full tissue field for each sample from which zoomed-in regions are shown in Fig. 1P main text). (C) Replicate 2 time-lapse series showing cell area extension (%) between consecutive stages, mapped on second time point. Both replicates show a similar distribution of growth between fruit tissues and stages. Abbreviations: v, valve; vm, valve margin; r, replum. Scale bars are indicated as 50 μm or 100 μm on each image. Note that the heat map for each sample scales the data between the maximum and minimum values in that sample. Therefore, it is important to consider the individual heatmap scales when comparing between samples.

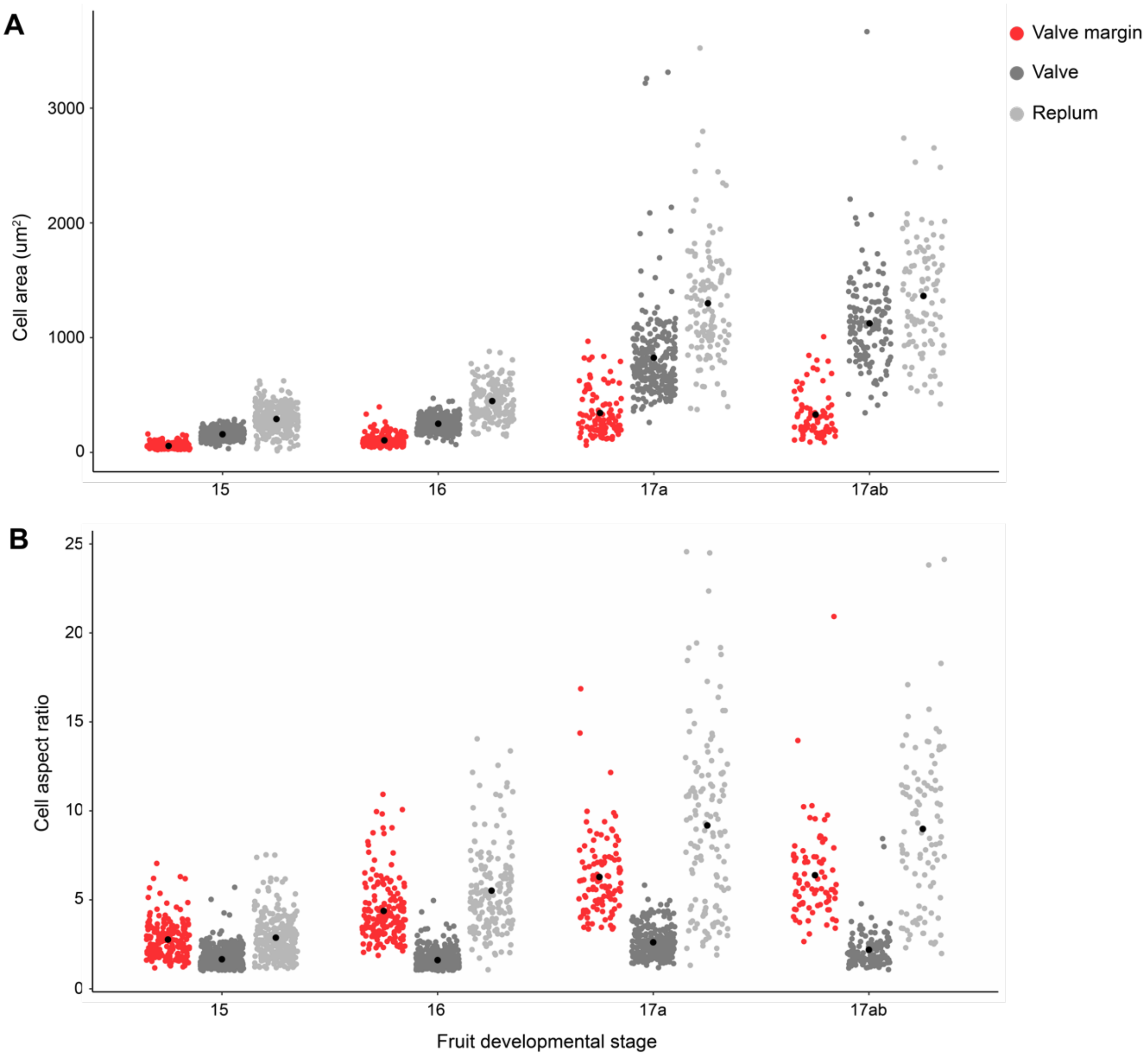


**Fig. S2 Valve margin cells are small and thin in *C. hirsuta* fruit. (A-B)** Cell area ($\mu m^2$) (A) and cell aspect ratio (length/width) (B) of cells in the valve margin (red), valve (dark grey) and replum (light grey) during stages 15, 16, 17a and 17ab of *C. hirsuta* fruit development; black dots indicate means; n = 2392 cells. Mean cell area of valve margin cells was significantly lower than that of valve and replum cells at all fruit stages ($P < 0.001$) based on two-way ANOVA (Tukey's HSD).

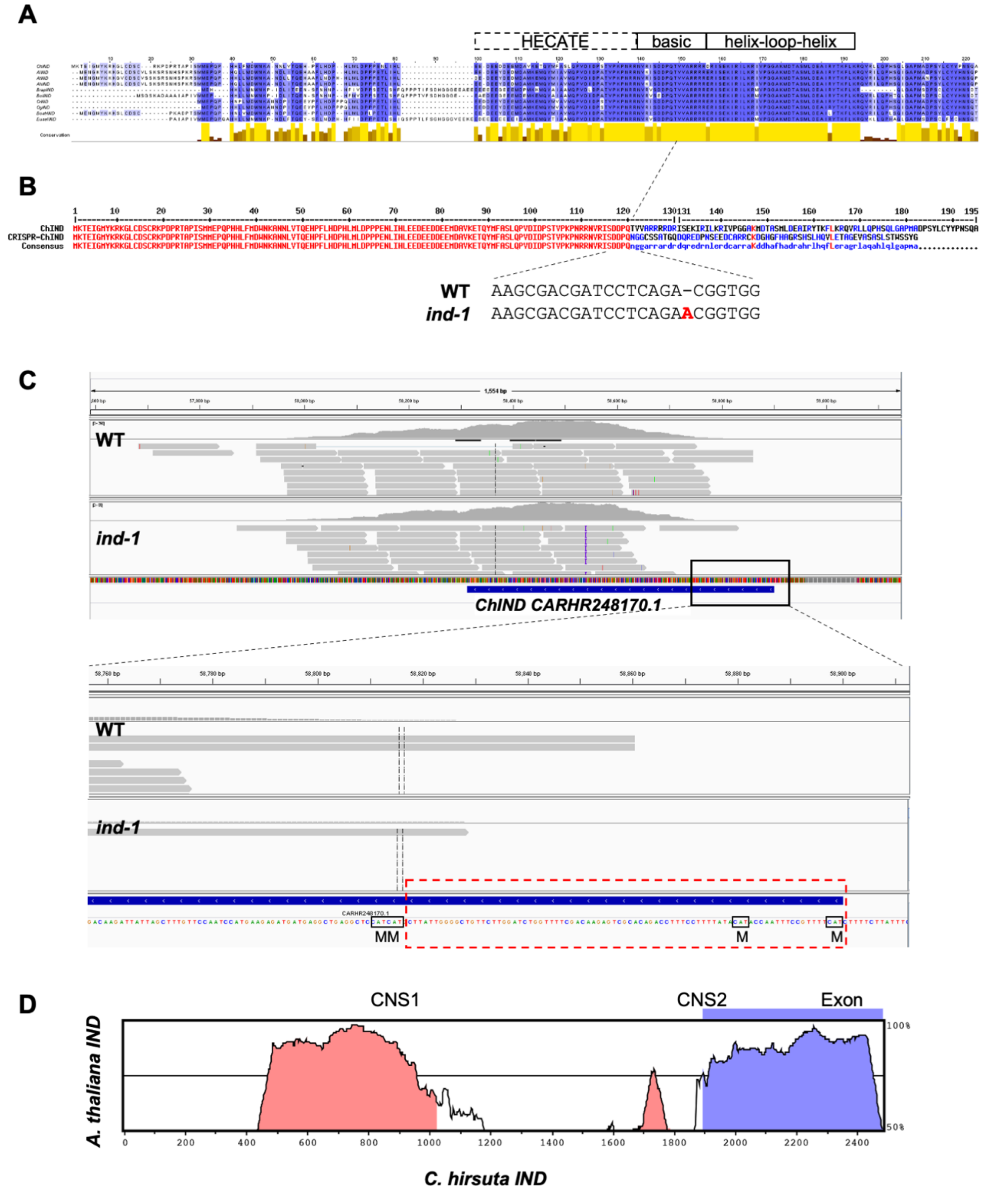


**Fig. S3 *C. hirsuta IND* sequence analysis.** (A) MUSCLE alignment (Waterhouse *et al.*, 2009) of IND amino acid sequences from 10 Brassicaceae species retrieved from Phytozome (Goodstein *et al*., 2012): *Cardamine hirstua, Arabidopsis lyrata, Arabidopsis thaliana, Arabidopsis halleri, Brassica rapa, Brassica oleracea capitata, Capsella rubella, Capsella grandiflora, Boechera stricta, Eutrema salsugineum*. Color code indicates percentage identity (blue ranges) and conservation percentage (yellow scale). Protein domains, indicated above alignment, were assigned according to (Pabon-Mora et al., 2014). Note that predicted protein sequences from Phytozome are shown here, but the starting methionine for *C. hirsuta* IND is likely to be the conserved methionine at position 32 in the alignment (Liljegren *et al*., 2004). (B) *C. hirsuta* IND amino acid sequence alignment with the *ind-1* allele produced by CRISPR/Cas9 editing. The single A insertion at position 361 bp of the coding sequence (shown in red) introduces a frame shift at amino acid position 121 and a premature stop codon, producing a truncated protein of 182 amino acids with 62 amino acids of nonsense protein sequence at the C-terminus. Red color in alignment indicates ≥ 90% consensus, blue color indicates <30% consensus. Note that the predicted protein sequence is shown here, but the starting methionine is likely to be M29 (Liljegren *et al.*, 2004). (C) Integrative Genomics Viewer screenshot of *C. hirsuta IND* (CARHR248170.1) gene together with RNA-seq reads from *C. hirsuta* wild type and *ind-1* mutant. Purple "I" in *ind-1* track indicates the single A insertion. A close-up view of the 5' gene region is shown below with four predicted methionine codons boxed and indicated by "M". Given that sequence reads map to an 84 bp region upstream of the putative start site at M29 (Liljegren *et al.*, 2004) (dashed red box), we distinguish this putative 5'UTR sequence from non-transcribed promoter sequence. (D) VISTA plot (Frazer *et al.*, 2004) showing 2480 bp of *C. hirsuta IND* (x-axis in bp) aligned with 2554 bp of *A. thaliana IND*; y-axis shows the percentage identity in 100 bp windows of the alignment. These sequences contain the single *IND* exon (blue) and whole 5' intergenic region of each gene. Two conserved non-coding sequences (CNS, pink) are identified; CNS1 includes the 406 bp valve margin element identified in *A. thaliana IND* (Girin *et al*., 2010).

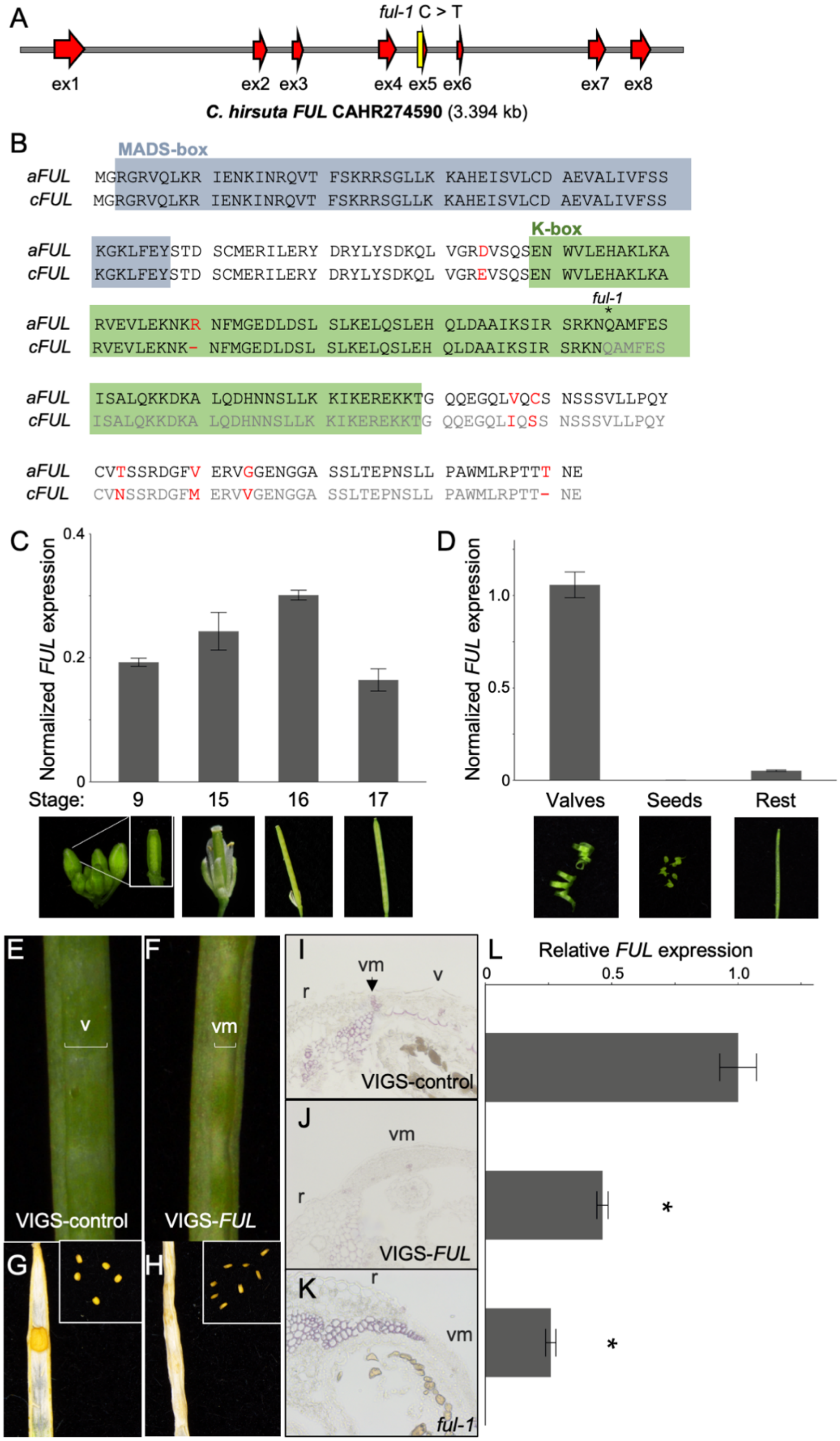


**Fig. S4** ***C. hirsuta FRUITFULL*** **(*FUL*).** (A) *C. hirsuta FUL* gene model with 8 exons (red arrows) and *ful-1* C to T mutation (yellow). (B) Alignment of FUL amino acid sequences from *A. thaliana* (*aFUL*) and *C. hirsuta* (*cFUL*), differences marked in red. MADS-box (grey) and K-box (green) domains shown according to the "Agamous-like MADS-box protein AGL8" entry in the UniProt Protein Knowledgebase. Premature stop codon at position 144 in *ful-1* is annotated '*' and greyed out amino acids are missing from the truncated FUL protein in *ful-1* mutants. (C-D) *C. hirsuta FUL* expression in wild-type fruit at stages 9, 15, 16 and 17 (C), and valve, seeds and remaining fruit tissues (rest, D), determined by quantitative RT-PCR and expressed relative to expression of the housekeeping gene *CLATHRIN*, bars represent mean ±SEM of three biological replicates. Photographs of sample types shown below. **(**E-L) VIGS of *C. hirsuta FUL.* Mature fruit (E-F), dry fruit and seeds (inset, G-H) and stage 17a phloroglucinol-stained transverse fruit sections (I-J) of VIGS-control (E, G, I) and VIGS-*FUL* (F, H, J); lignified cell walls stain pink. Stage 17a phloroglucinol-stained transverse fruit section of *ful-1* shown as control (K). *C. hirsuta FUL* expression in VIGS-control, VIGS-*FUL* and *ful-1* 17a stage fruit, determined by quantitative RT-PCR, normalized to expression of the housekeeping gene *CLATHRIN*, and shown as relative to VIGS-control, bars represent mean ±SEM of technical replicates of a single fruit. Asterisks indicate statistically significant differences between means of technical replicates ($P < 0.05$; Student's t-test) (L). VIGS fruit were divided in half and one half was used for phloroglucinol staining and the other half for quantitative RT-PCR.

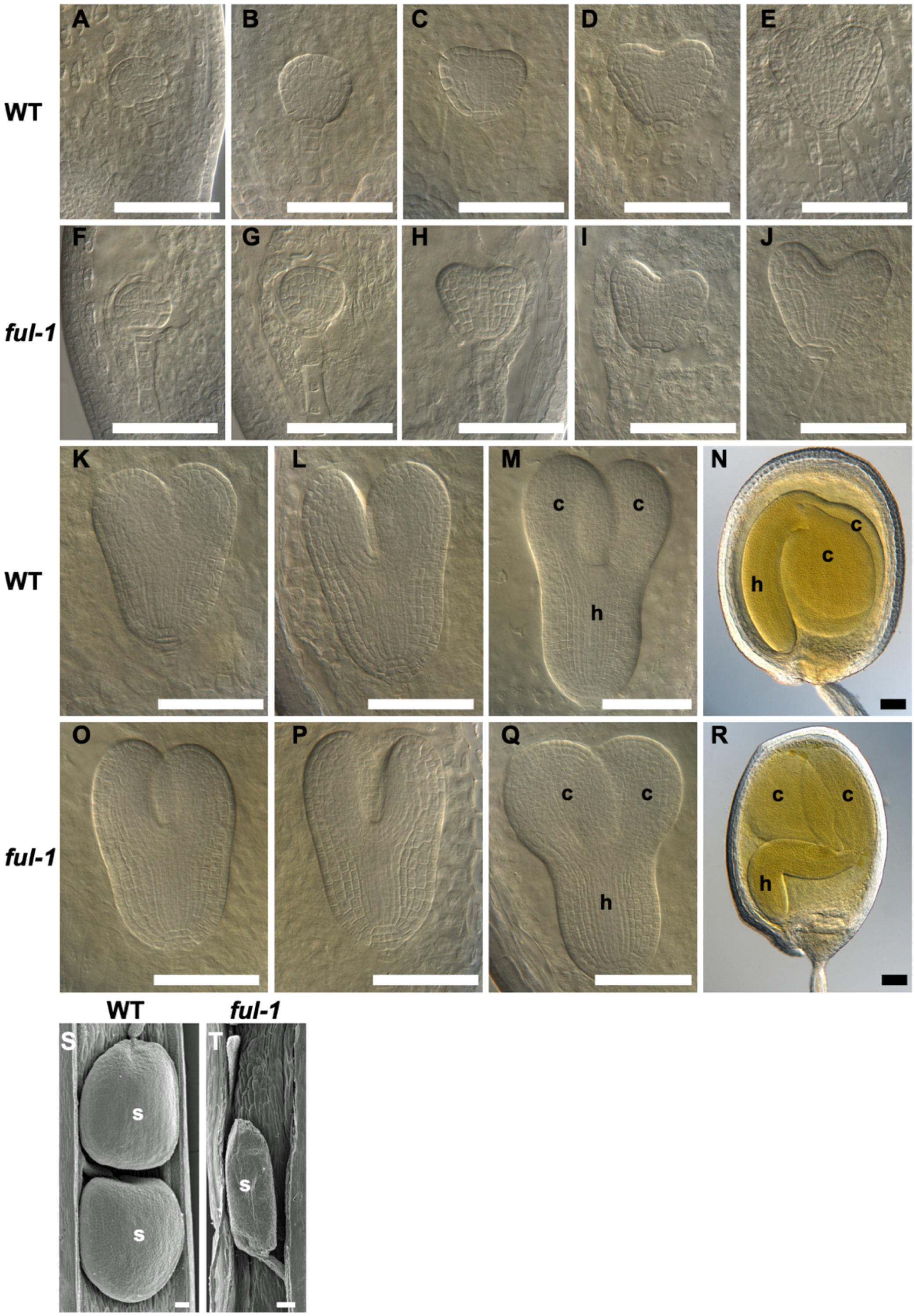


**Fig. S5 Embryo development in *C. hirsuta* wild-type and *ful-1* seeds.** (A-R) Differential interference contrast micrographs of embryos at globular (A, B, F, G), heart (C-E, H-J, K, L, O, P), torpedo (M, Q) and bent-cotyledon (N, R) stages. (S-T) SEMs of seeds in stage 17b fruit. Genotypes indicated. Abbreviations: h, hypocotyl; c, cotyledon. Scale bar: 100 μm.

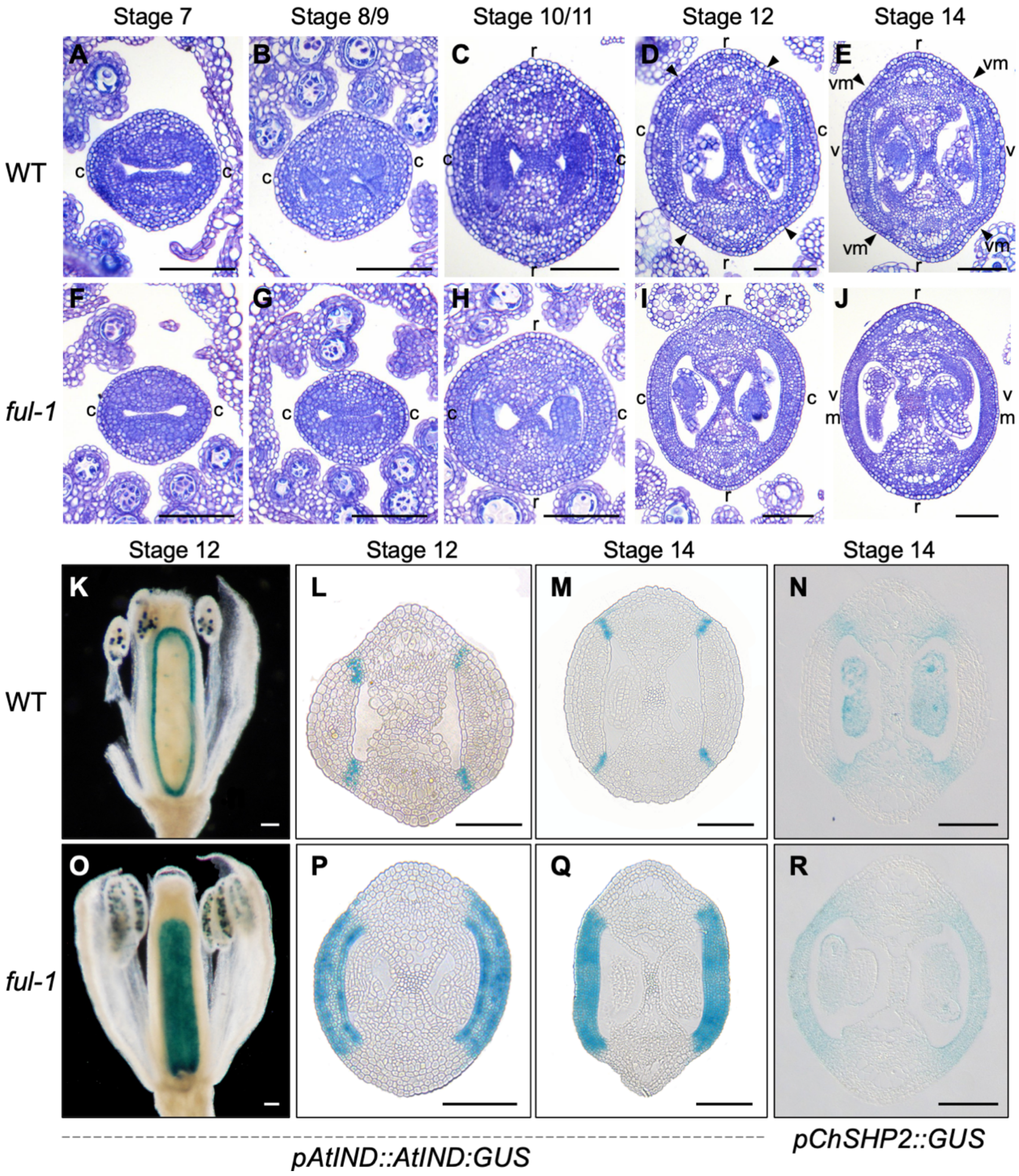


**Fig. S6 Gynoecium and early fruit development and expression of valve margin identity genes in *C. hirsuta* wild type and *ful-1*.** (A-J) TBO-stained transverse sections of *C. hirsuta* wild type (A-E) and *ful-1* (F-J) during gynecium and early fruit development. Developmental stages indicated above panels. Arrows mark boundary between replum and carpel where the valve margin forms. (K-R) GUS-stained fruit of *C. hirsuta* wild type (K-N) and *ful-1* (O-R). *AtIND::IND:GUS* expression (blue) marks tissues with valve margin identity in whole mount (K, O) and transverse sections (L-M, P-Q). *ChSHP2::GUS* expression (blue) marks tissues with valve margin identity in transverse sections (N, R). Developmental stages indicated above panels. Abbreviations: c, carpel; r, replum; v, valve; vm, valve margin. Scale bars: 100 µm (A-J, L-N, P-Q), 1 mm, (K, O).

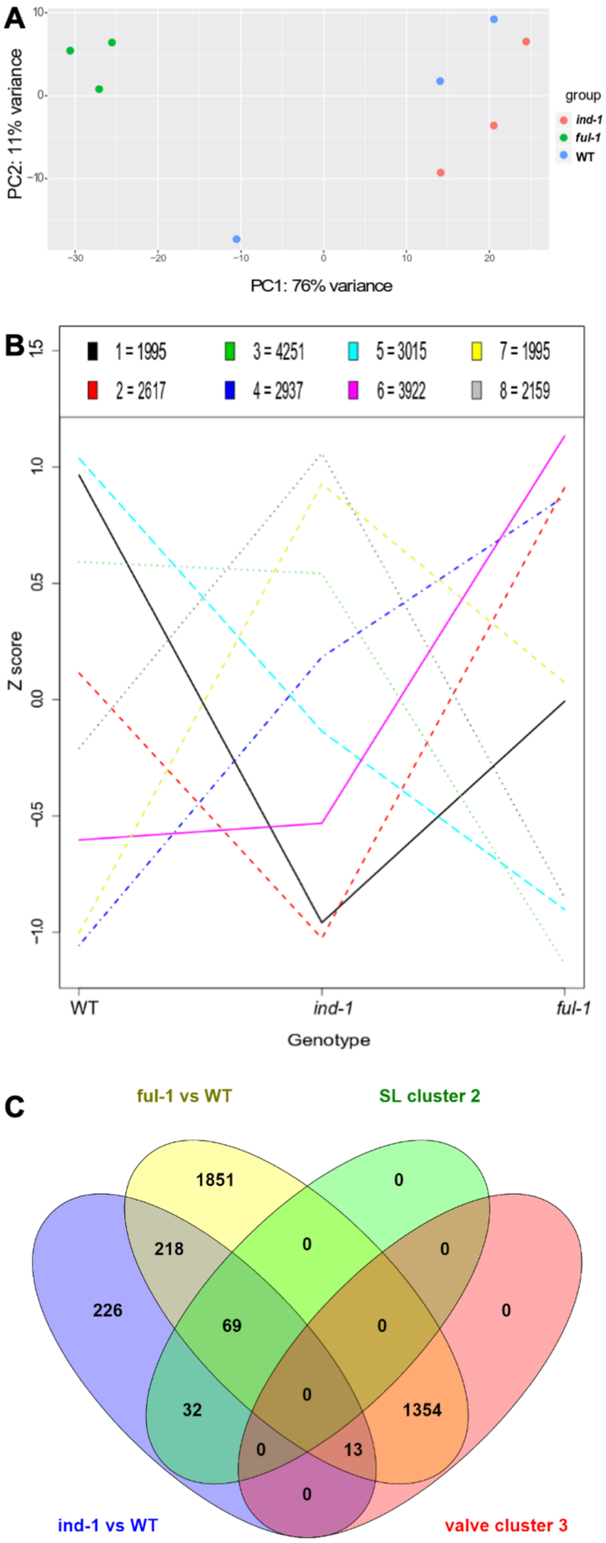


**Fig. S7 RNAseq analysis of *C. hirsuta* wild-type, *ind-1* and *ful-1* stage 17ab fruit.** (A) Principal component analysis of all RNAseq samples indicates that replicates cluster according to genotype, except for one outlier wild-type sample (blue) that was removed from further analysis. (B) Plot of eight gene expression clusters with number of genes defining each cluster shown in key. Clusters 2 and 3 are shown in main Fig. 4K. (C) Venn diagram comparing the genes in cluster 2 (separation layer-associated) and cluster 3 (valve-associated) with the differentially expressed genes between *ind-1* and wild-type fruit, and *ful-1* and wild-type fruit.

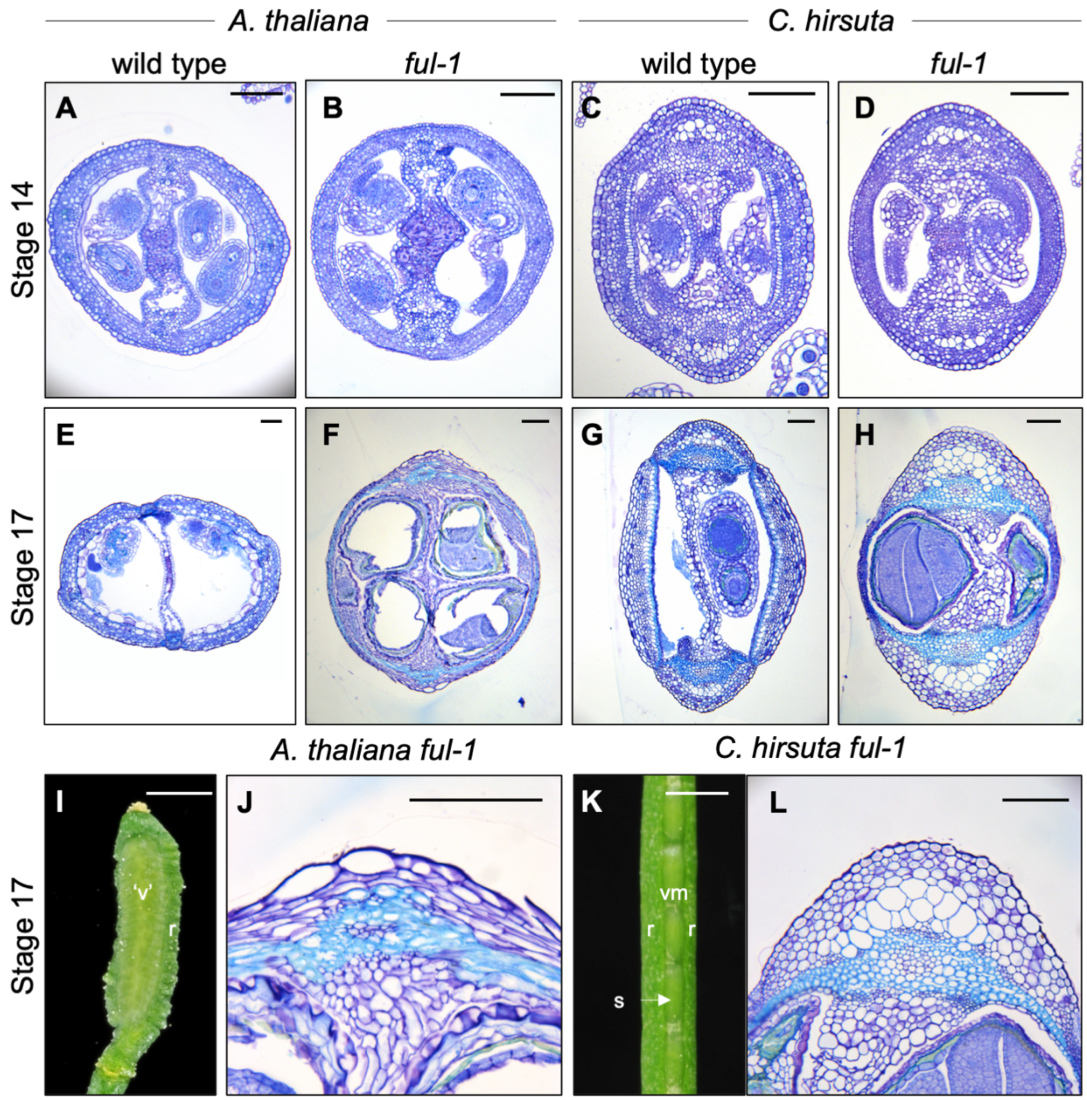


**Fig. S8 Histology of *A. thaliana* and *C. hirsuta* wild-type and *ful-1* fruit.** (A-H) TBO-stained transverse sections of stage 14 (A-D) and 17 (E-H) fruit of *A. thaliana* wild type (A, E), *ful-1* (B, F), and *C. hirsuta* wild type (C, G), *ful-1* (D, H). (I-L) Stage 17 fruit of *A. thaliana ful-1* (I, J) and *C. hirsuta ful-1* (K, L). Close-up views of (F, H) are shown in (J, L). Abbreviations: v, valve; r, replum; vm, valve margin; s, seed; 'v', valve-like tissue. Scale bars: 100 μm (A-H, J, L), 1 mm (I, K).

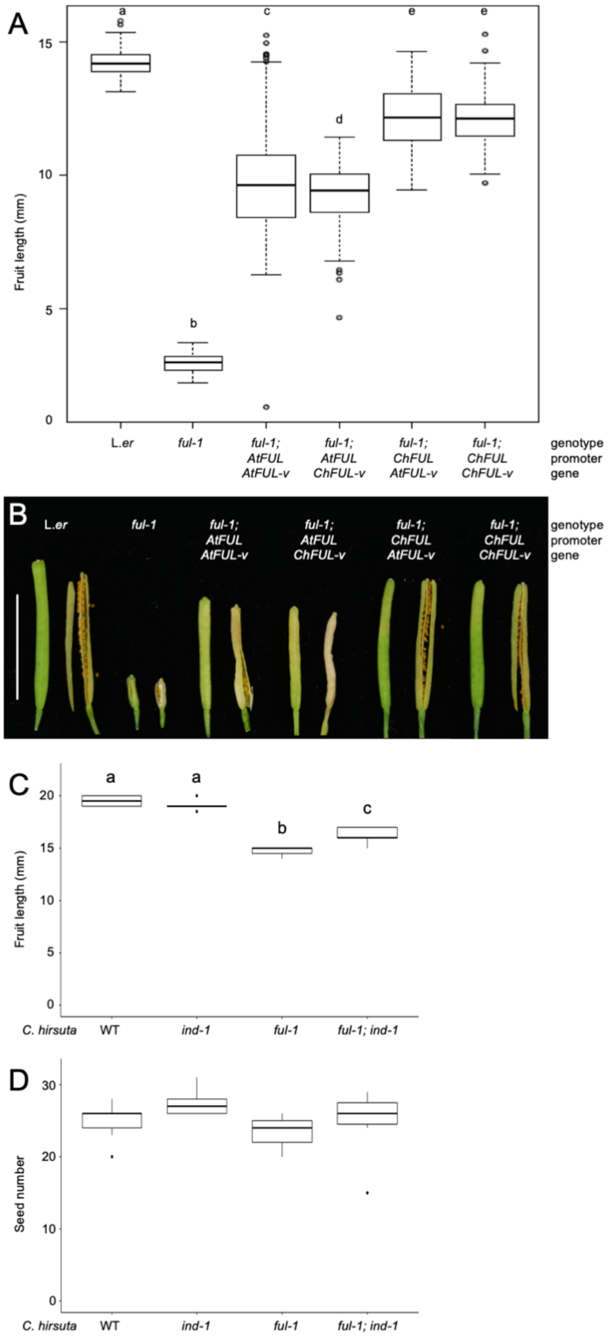


**Fig. S9: Complementation of *A. thaliana ful-1* with *C. hirsuta FUL*.** Fruit length (A) and dehiscence (B) in *A. thaliana* Landsberg *erecta* wild type, *ful-1*, and *ful-1* complemented with *pAtFUL::gAtFUL-Venus*, *pAtFUL::gChFUL-Venus*, *pChFUL::gAtFUL-Venus*, and *pChFUL::gChFUL-Venus*, n = approximately 20 fruit from 3-15 plants for each genotype including 4 independent *ful-1; pChFUL::gChFUL-Venus* T2 lines. Fruit length (C) and seed number (D) in *C. hirsuta* wild type, *ind-1*, *ful-1*, *ind-1; ful-1*, n = at least 5 fruit for each genotype. Letters denote statistically significant differences (P < 0.05) between means based on one-way ANOVA (Tukey's HSD). Note that no significant differences were found between samples in (D).

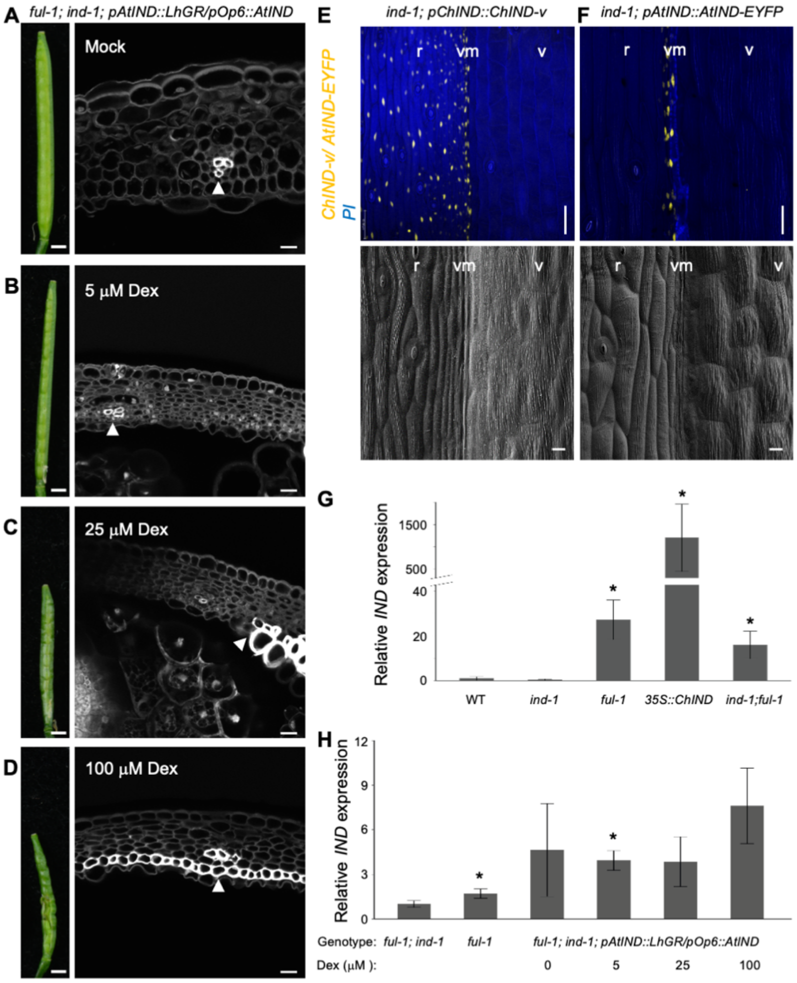


**Fig. S10: Ectopic *IND* dosage contributes to whether valve margin cells adopt lignified or separation layer fate.** (A-D) Dosage series of mock (A), 5 µM (B), 25 µM (C) and 100 µM (D) dexamethasone applied to *ful-1; ind-1; pAtIND::LhGR/pOp6::AtIND* plants. Representative images of fruit (left) and CLSMs of valve transverse sections stained with PI to outline cells (right); arrows indicate lignified cell walls. Homozygous plants from a T3 line with a single transgene insertion were used. Dexamethasone was applied repeatedly over a 2-week period during which the fruit analyzed developed through to stage 17b. Compared to the valves of mock-treated *ful-1; ind-1 AtIND>GR>IND* plants (A), Dex-treated valves contain small valve margin cells with presumed separation layer identity; an increasing number of cells with lignified layer identity developed in response to increasing *IND* dose (B-D). (E-F) CLSM and SEM of stage 17 *C. hirsuta ind-1* fruit expressing *pChIND::ChIND-v* (E) or *pAtIND::AtIND-EYFP* (F); IND expression (yellow), PI stain (blue). Complementation of valve margin cells and functional dehiscence zones is observed in both transgenic lines, despite the more restricted expression domain of *A. thaliana IND* compared to *C. hirsuta IND*. (G-H) *C. hirsuta IND* expression in stage 17 fruit of wild-type (wt) compared to *ind-1*, *ful-1*, *35S::ChIND* and *ful-1; ind-1*, (G) and *ful-1; ind-1 AtIND>GR>IND* mock-treated fruit compared to fruit treated with dexamethasone concentrations of 5 μM, 25 μM and 100 μM (H), determined by quantitative RT-PCR, normalized to expression of the housekeeping gene *CLATHRIN*, and expressed relative to wild type (G) or *ful-1; ind-1* (H). Homozygous plants from a T3 line with a single transgene insertion were used in (H). Error bars represent standard error of two (*ful-1; ind-1 AtIND>GR>IND*) or 3 (wt, *ind-1*, *ful-1*, *35S::ChIND* and *ful-1; ind-1*) biological replicates. Asterisks indicate statistically significant differences between means of biological replicates ($P < 0.05$; Student's t-test) compared to wild type (G) or *ful-1; ind-1* (H). Abbreviations: v, valve; r, replum; vm, valve margin, Dex, dexamethasone, PI, propidium iodide. Scale bars: 1 mm (fruit, A-D); 10μM (CLSMs, A-D and SEMs, E-F); 50μM (CLSMs, E-F).
